# Predicting the immediate and subsequent effects of commercials on product valuation using EEG and deep learning

**DOI:** 10.64898/2026.09.22.751523

**Authors:** Inbal Gur Arie, Daniel Andrew Atad, Adam Hakim, Dino Levy

## Abstract

Neuromarketing mainly seeks to enhance the prediction of marketing stimuli success, such as commercials and movie trailers, by integrating neurophysiological measures with traditional behavioral measures. In the current study, the authors tested whether neural activity could predict consumer valuation and how preferences change after watching commercials, both immediately and over time. Participants (n=161) watched images of products followed by commercials advertising those products while their neural activity was recorded using electroencephalograph (EEG). They watched the product images again a week later without EEG recordings. Immediately after each stimulus exposure, participants stated their willingness to pay (WTP) for the product and how much they liked the commercial. Standard EEG measures showed weak and inconsistent relationships with behavior and yielded near-chance predictions. In contrast, deep learning models applied to the raw neural data achieved substantially higher predictive accuracy, successfully predicting both immediate and delayed WTP and ad liking. Importantly, the authors were able to successfully predict preferences for new participants and/or new products the models were not trained on. However, accuracy declined as generalization demands increased. Moreover, prediction performance declined as value differences narrow. Together, these findings show that neural responses carry reliable and temporally persistent information about consumer valuation and its evolution over time.

## Introduction

Neuromarketing seeks to enhance the prediction of marketing stimuli success, such as commercials and movie trailers, by integrating neurophysiological measures with traditional behavioral measures. While behavioral measures, such as questionnaires, focus groups, and interviews are the primary means of prediction in the industry (McDaniel & Gates, 2016; Wilkinson & Birmingham, 2003), their effectiveness is limited (Karmarkar & Yoon, 2016; Smidts et al., 2014). Behavioral measures face limitations due to self-report biases and subconscious reactions (Calvert & Brammer, 2012), leading to inconsistencies in predicting outcomes (Fisher, 1993; Johansson et al., 2006; Mcdaniel et al., 1985; Neeley & Cronley, 2004) and have high failure rates, ranging from 40% to 80% (Castellion & Markham, 2013).

It has been shown that using neural activity can increase the prediction success above and beyond behavioral measures (Boksem & Smidts, 2015; Christoforou et al., 2017; Costa-Feito et al., 2023; Genevsky et al., 2017; Hakim et al., 2018, 2021; Karmarkar & Plassmann, 2017; Motoki et al., 2020; Venkatraman et al., 2015) suggesting that using neural activity for prediction can overcome some of the biases and problems that behavioral measures have.

Despite advancements in the field, significant gaps persist. No EEG studies have examined if we can use neural activity to predict if and to what extent consumers’ preferences for an advertised product change after watching a commercial for that product. Successful prediction of commercial’s effect on product valuation is an important goal in the industry as it provides an objective an unbiased prediction metric. In the current study we aim to fill in this gap.

The present study advances prior research in four systematic ways. First, we address the pervasive limitation of small sample sizes by collecting EEG data from 161 participants, representing one of the largest datasets in this domain. The scale of the dataset enables us to implement and compare a range of deep learning models, thereby assessing the robustness of neural prediction across different modeling approaches. Second, we extend existing approaches that focus predominantly on single-time-point predictions by examining how exposure to commercials influences product valuation across two temporal horizons: immediately following exposure, and after a one-week delay. Third, we expand the scope of prediction to encompass not only shifts in product valuation but also preferences for the commercials themselves, thereby providing a more comprehensive account of how advertising stimuli shape consumer decision-making. Fourth, we compare prediction accuracies across several out-of-sample training and testing procedures. From procedures that use all stimuli and participants for training, towards the more demanding but more ecological approaches of predicting preferences for participant and/or products the models were not trained on. By replicating our earlier findings (Hakim et al., 2023), while systematically extending them along these dimensions, this study delivers a robust and generalizable test of the capacity of neural signals to predict changes in consumer valuation.

In addition, we also integrate and compare two major methodological approaches in the field. First, we implement the traditional feature-based framework, extracting well-established EEG markers such as spectral band power, inter-subject correlation (ISC), and frontal asymmetry (FA), and modeling these predictors using logistic and linear regressions. This approach reflects the dominant strategy in most prior work and provides a benchmark for assessing interpretable neural contributions to consumer decision-making. Second, we use feature-free deep learning approach that operates directly on the neural data, inspired by recent evidence showing that deep learning models can outperform feature-engineered pipelines in predicting consumer preferences (Hakim et al., 2023). By evaluating both approaches on identical prediction targets, we can quantify the incremental value of end-to-end neural network models relative to classical feature-based methods. Anticipating our results, the deep learning models consistently exhibited superior predictive accuracy, highlighting the potential of advanced techniques to capture subtle neural signatures of valuation that traditional features may overlook.

## LITERATURE REVIEW

Integrating neurophysiological measurements, such as EEG and functional magnetic resonance imaging (fMRI), alongside behavioral measurements, has provided deeper insights into consumer behavior by measuring real-time brain activity in response to marketing stimuli (Agarwal & Dutta, 2015; H.-Y. Chan et al., 2024; Gupta et al., 2025; Harris et al., 2018; Hsu & Yoon, 2015; Knutson & Genevsky, 2018; Lin et al., 2018; Murugappan et al., 2014; Newton-Fenner et al., 2023; Smidts et al., 2014). The scope of these techniques is broad, and can be used to investigate different aspects of marketing research (Khurana et al., 2021) like brand evaluation (Esch et al., 2012), brand preferences (Yu et al., 2018) and market success (Baldo et al., 2015; Barnett & Cerf, 2017; Boksem et al., 2025; Boksem & Smidts, 2015), among others. This breadth demonstrates the potential of neurophysiological methods to reveal underlying cognitive and affective processes that shape consumer behavior, laying the foundation for their use in predictive applications.

Prediction has become a central focus in neuromarketing research. At the individual level, studies used EEG to predict consumer choices in natural environments (Horr et al., 2022) and to estimate the likelihood of product purchase (Hakim et al., 2023; Pratama et al., 2024). Extending beyond individual decisions, neural activity from relatively small laboratory samples has also been linked to broader market outcomes, such as anticipating product satisfaction (Kumar et al., 2019), the market success of consumer goods (Berns & Moore, 2012; Genevsky et al., 2025; Knutson & Genevsky, 2018), and large-scale behaviors such as YouTube viewership (Falk et al., 2012; Hakim et al., 2021; Knutson & Genevsky, 2018), retail sales (Kühn et al., 2016; Varga et al., 2021; Venkatraman et al., 2015), crowdfunding (Genevsky et al., 2017, 2025), and article sharing (Scholz et al., 2017). Collectively, this literature demonstrates that neural data can improve predictive accuracy across multiple domains, including product\ad preferences and willingness to pay. Within this broad literature, commercials represent a particularly central and ecologically valid testbed for examining how neural responses during ad viewing can be used for prediction.

Building on this central role, a growing body of neuromarketing research has applied neurophysiological and physiological tools, primarily EEG and fMRI, to assess how commercials affect viewers at both individual and population levels (H.-Y. Chan et al., 2024; Guixeres et al., 2017; Motoki et al., 2020; Ohme et al., 2009; Venkatraman et al., 2015). EEG has been extensively used to decode neural correlates of ad effectiveness. Studies show that neural synchrony across viewers, quantified through inter-subject correlation, can predict commercial success by indexing shared attentional and emotional engagement (Barnett & Cerf, 2017; Dmochowski et al., 2014). Elevated ISC has been linked to improved recall, stronger emotional resonance, and increased likelihood of ad virality or product choice. Additional EEG metrics such as frontal alpha asymmetry (reflecting approach motivation), enhanced theta and gamma activity (associated with narrative engagement and emotional arousal), and ERP components like the P300 (linked to attentional allocation) have also been found to track reactions to advertisements (Dimpfel, 2015; Guixeres et al., 2017; Vecchiato et al., 2011, 2014; Wang et al., 2016). While these studies highlight the promise of traditional neural metrics for explaining ad effectiveness, moving beyond these approaches toward machine learning and deep learning is essential for systematically improving prediction – a direction the present study undertakes.

Recent advances in machine and deep learning have already begun to transform neuromarketing research, with several studies demonstrating that these methods can extract predictive patterns from EEG and fMRI data that traditional neural metrics alone cannot capture (Aldayel et al., 2020; Byrne et al., 2022; H.-Y. Chan et al., 2024; Hakim et al., 2021, 2023; Ishtiaque et al., 2025; Motoki et al., 2020; Xu & Liu, 2024; Zeng et al., 2022; Zhao et al., 2025). For example, Hakim et al., (2021) used machine learning on EEG recordings collected during ad viewing and found that brain signals could outperform traditional questionnaires in predicting which commercials participants preferred and how those ads influenced product valuation. Other studies have reached similar predictive insights. Motoki et al., (2020) showed that neural activity in key social-cognition regions forecasted the likelihood of a commercial being shared on social media beyond what participants’ self-reported intentions would predict. Similarly, Chan et al., (2024) found that early neural responses associated with emotion and memory formation were strong indicators of how much viewers liked an advertisement, while activity in mentalizing regions (involved in understanding others’ thoughts) better predicted whether an ad would go “viral” on a larger scale.

These findings underscore the promise of machine learning for predicting advertising impact, among other outcomes. However, applications remain constrained by small sample sizes, and most studies have not implemented machine or deep learning – based predictive approaches on the raw EEG data, continuing instead to rely primarily on traditional neural metrics such as specific ERP components (e.g. N200, P300), frequency band power (e.g. alpha, beta, theta), or frontal asymmetry (Bilucaglia et al., 2025) – limitations that motivated the methodologies adopted in the present study.

Building on this foundation, the present study advances the field by combining EEG with deep learning models in a uniquely large dataset of 161 participants, enabling the systematic comparison of multiple predictive models. In doing so, we replicate our prior findings (Hakim et al., 2023) and extend them by predicting product valuation across three assessment points – after viewing the product image (pre-ad baseline), immediately after viewing its commercial, and again one week later. Moreover, we also predict the preferences for the commercials themselves. Hence, our study provides a systematic and large-scale demonstration of how neural signals can be used to predict willingness to pay and consumer preferences at multiple time points. Finaly, we show that we can predict valuation for new products and/or for new consumers that the models were not trained on. In our opinion, this out-of-sample prediction demonstrates what the industry cares about – generalizability and usability in real world market research scenarios.

## METHODS

### Participants

161 participants took part in the study (69 women, ages 20-63, mean = 27.04±7.36). All participants gave written informed consent before participating in the study, which was approved by the local ethics committee at our university. Out of all participants, 24 participants participated for school credit and the remaining participants received $20 show up fee. We omitted the data of 17 participants due to technical problems of the EEG recordings and additional 6 participants that did not meet our exclusion criterion (see below).

### Procedure

See figure 1 for a detailed description of the study design. There were three parts to the experiment. The first two parts were conducted consecutively in the lab on the same day, while we recorded participants’ brain activity using an 8-electrode EEG device (Neuroelectrics, Spain; see section 2.5). The third part was conducted online, one week later without EEG recordings.

**Figure 1.**
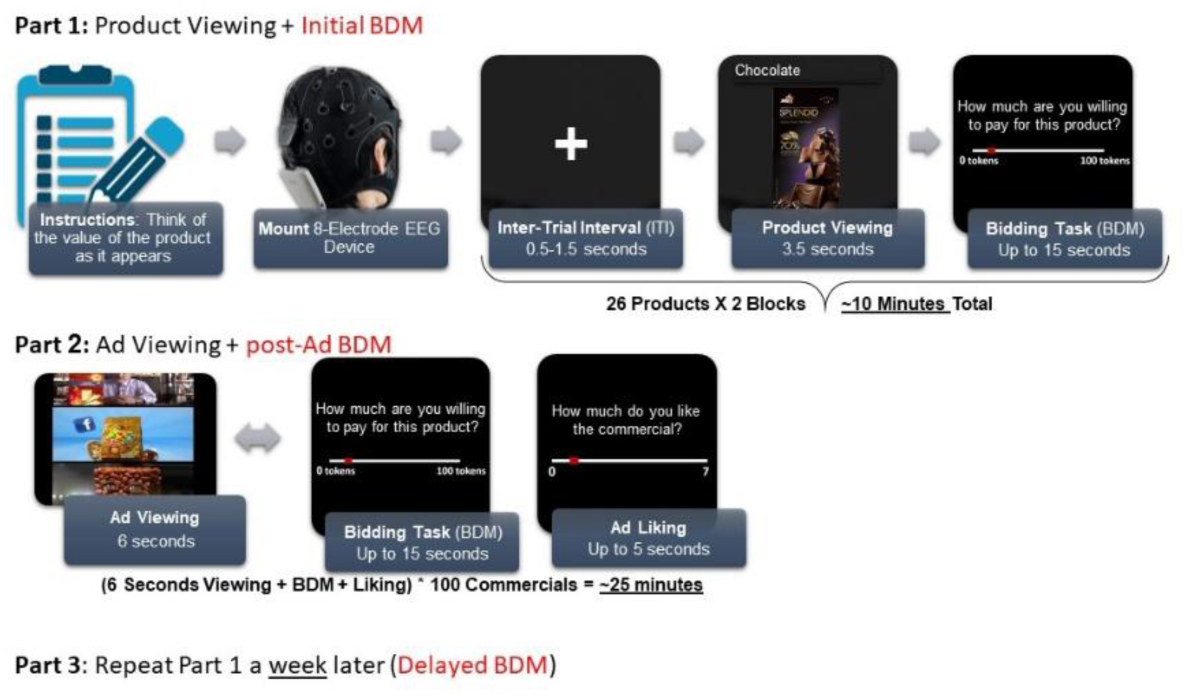
Study design. Participants completed three parts of the experiment, depicted as Part 1, 2 and 3. In Part 1, on each trial, participants viewed a picture of a consumer product and indicated their WTP for the product. In Part 2, participants watched a commercial of a consumer product and indicated their WTP for the product appearing in the commercial and overall liking for the commercial. During these two parts, participants’ EEG signal was recorded. Part 3 was identical to Part 1, though without EEG recording and conducted online outside the lab. Note, that the products advertised in the commercials were the same products presented in Parts 1 and 3.

#### Part 1

During the first part of the experiment, on each trial, participants viewed a picture of a consumer product for 3.5 seconds. Above the picture, there was a short description of the product (two to three words). Afterwards, a slider between zero and a hundred tokens appeared on the screen. Above the slider, the question “how much are you willing to pay for this product?” was presented. Participants stated their WTP for the product by moving the cursor of the slider using the computer mouse. On each trial, we randomly positioned the cursor such that participants could not anticipate if, how far, and to which direction, they will need to move the cursor. This allowed us to avoid motor preparation neural signals which could be correlated with value. The slider appeared until a response, with a maximum time of 15 seconds, after which the trial was over. We had 26 products, each of them was presented twice in a randomized order. One product (Baracke) was introduced later in the data collection and was therefore not available for a subset of participants (n = 41); nevertheless, it was retained in all analyses.

#### Part 2

In the second part of the experiment, which started immediately after the first part, on each trial, participants viewed a commercial of a consumer product for 6 seconds. We presented, for each product, three to five different commercials, for a total of one hundred commercials. All commercials advertised the same 26 products that were presented in Part 1 (including one product that was not available for a subset of participants; see Part 1). After each commercial, two questions appeared one after the other (in a counterbalanced order). The first question was identical to the question from the first part – “how much are you willing to pay for this product?”. Again, participants indicated their WTP on a slider between zero and a hundred tokens. The second question was “how much did you like the commercial that was just presented?” on a continuous slider between zero and seven. Participants had maximum 15 seconds to answer each of the questions. Here also, the location of the cursor was randomly positioned for each question.

#### Part 3

A week later, participants completed the third part of the experiment online. In this part, exactly as in Part 1, they had to indicate how much they are willing to pay for the same products they previously saw on Parts 1 and 2. Here, again, each product was presented twice in a randomized order. They completed the experiment using Qualtrics after receiving the experiment link via an email.

### BDM and Willingness to Pay

To make the experiment incentive compatible, we used the standard BDM procedure (Becker et al., 1964) in order to estimate participants’ willingness to pay for each of the products they saw. We conducted this procedure twice – once after the lab session (Parts 1 and 2) and once after the online session. After completing the experiment, the computer randomly selected one trial from Parts 1 and 2 and another from Part 3, meaning that two different products could potentially be won. As a result, some participants won no products, some won one, and others won two. The products appearing in these trials were assigned a random price between 0 and 100 tokens by the computer. If the amount participants offered for the product in this specific trial exceeded the price generated by the computer, participants received the product at the random price and kept the remaining change. If participants offered less than the random price, they did not win the product and received only the change left. We converted the 100 tokens to ₪10. Thus, for example, if a participant won a product for 52 tokens in the first random trial, and did not win a product in the second random trial, her total winnings were ₪80 participation fee, the product, and ₪4.8 from the first trial (she is left with 48 tokens change that is, ₪4.8), and ₪10 from the second trial, summing to a total of ₪94.8.

### Stimuli

In Part 1, there were 26 different products from five categories: beverages, breakfast, dry goods, hygiene, and snacks. See table w1 in the web appendix for the full list of the products and figure 2 for a few examples.

**Figure 2.**
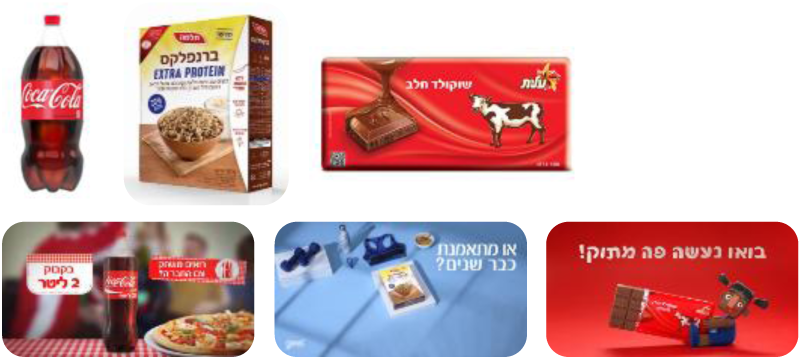
Examples of the stimuli. On the top, three pictures of different products – Coca Cola, Branflakes extra, and milk chocolate. On the bottom, frames from commercials of the same products.

### EEG system and recording procedure

We used an 8 wet electrode EEG device (StartStim 8, Neuroelectrics, Spain). The EEG signal was sampled at 500 Hz, with a bandwidth of 0–125 Hz (DC Coupled), resolution of 24 bits – 0.05 lV, measurement noise was below 1 lV RMS, common mode rejection ratio is 115 dB and input impedance was 1GX. We positioned the electrodes at positions F7, Fp1, Fpz, Fp2, F8, Fz, Cz, Pz (according to the standard 10-20 system). See figure 3 for a description of the electrode montage.

**Figure 3.**
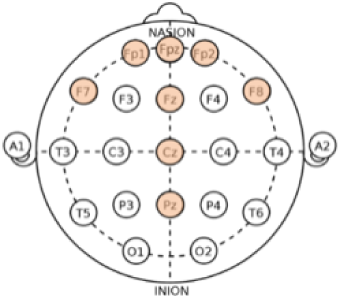
Electrode montage. Location of electrodes used are painted orange.

### Statistical analyses

#### Behavioral preprocessing

We normalized each participant’s data, both their WTP and liking values, to ensure all participants are on the same scale. We further wanted to make sure that participants’ bids were different across products so each participant will demonstrate clear preference ordering of the products. Therefore, we examined each participant’s bids. Figure 4 represents the BDM bids of three example participants in each of the three parts of the experiment. As can be seen, each of the participants demonstrated a clear preference ordering across the entire range of the products. We used an exclusion criterion based on each participant’s variance in their WTP values, such that participants with a standard deviation smaller than five were excluded as it indicated that they used only a very limited price range. Six participants were excluded based on this criterion.

**Figure 4.**
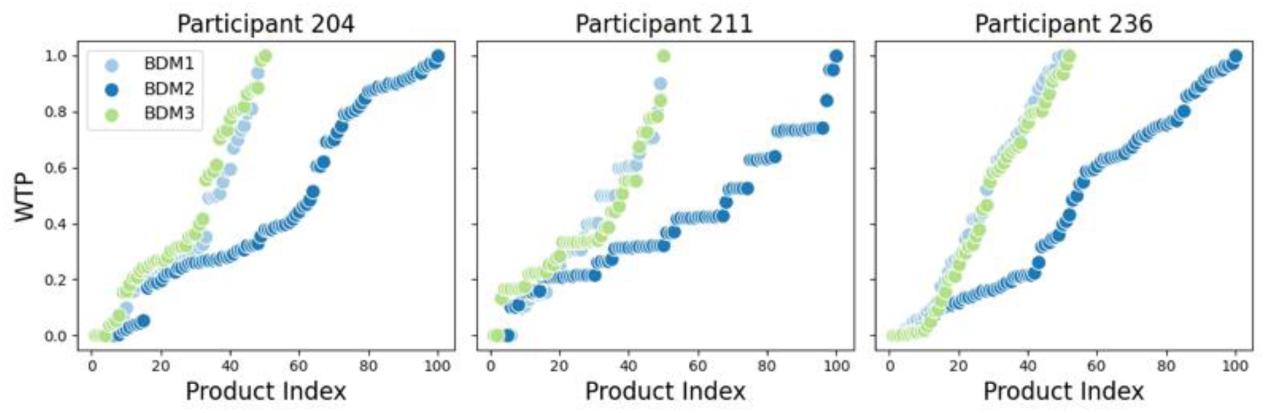
WTP distribution of 3 different participants. Part 1, Part 2, and Part 3 represent the WTP values for each product in each part, respectively. Trials are arranged in ascending order based on WTP values.

### EEG preprocessing

EEG preprocessing was conducted in line with the procedures reported by Hakim et al., (2021). EEG recordings were segmented to epochs such that each epoch consisted of the stimulus’ viewing period and 1 second of the previous ITI used as baseline activity. We did not use the response period to measure a value signal to avoid contamination by motor movements. The length of each epoch was 4.5 seconds in Part 1 and 7 seconds in Part 2, corresponding to 3.5 seconds and 6 seconds stimulus length, respectively. A sample, then, consisted of 8 channels and 2250 time points (4.5 seconds at 500Hz) for Part 1 and 3500 time points for Part 2 (7 seconds at 500Hz). Then, each epoch was cleaned using a bandpass filter between 0.5Hz and 100Hz and notch filter of 50Hz. Next, for each data sample, if any electrode recording showed a standard deviation greater than 50 mV or a maximum amplitude exceeding 400 mV, that electrode’s data was discarded and replaced with data from a neighboring electrode (Delorme & Makeig, 2004). However, if four or more electrodes in a sample met these exclusion criteria, the entire sample was discarded. For the deep learning analyses, data was further down-sampled to 250 Hz, in order to facilitate reasonable processing times.

### Behavioral analyses

#### WTP reliability

We wanted to make sure that we can use participants’ reported WTP values as stable representations of their product valuations for the prediction attempts. Thus, we checked for the correlation of the WTP values between the first and second time a product appeared in both Part 1 and Part 3. For the WTP values in Part 2, as each product repeated between three to five times, we conducted an intraclass correlation coefficient analysis.

### Commercials’ influence

We examined how the commercials influenced participants’ WTP and whether these effects persisted across time. To this end, we fit a linear mixed-effects model with Part (Part 1, Part 2, Part 3) as a fixed effect, random intercepts for participants and products, and WTP as the dependent variable. Part 1 served as the reference level, such that the fixed-effect coefficients estimated mean differences in WTP for Part 2 vs. Part 1 and Part 3 vs. Part 1. To fully assess differences between all sessions, we conducted planned pairwise comparisons (Part 2– Part 1, Part 3– Part 1, Part 2– Part 3) obtained from the fitted model with Holm correction to adjust for multiple testing. Holm correction was chosen because only three pre-specified comparisons were tested, and in such cases, it is preferable to control the overall probability of making any false positive error; by contrast, false discovery rate (FDR) correction was used in later analyses involving a larger number of exploratory tests. The fitted regression model was

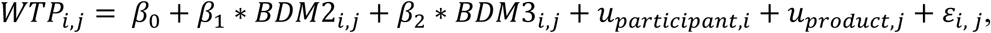

where *β*_0_ is the intercept (WTP in Part 1), *β*_1_ and *β*_2_ represent fixed effects of the second and third parts, *u_participant,i_* and *u_product,j_* are random intercepts for participant *i* and item *j*, and *ε_i_*, *_j_* is the residual error.

### WTP and liking relation

We wanted to examine if participant’s liking for a commercial is correlated with their WTP for the product appearing in the commercial. Therefore, we conducted a simple linear regression with clustered errors where WTP was the dependent variable and *Liking* was the predictor. The fitted regression model was:

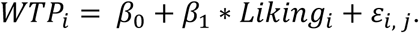

### EEG analyses

#### EEG spectral power

After the preprocessing stage, we performed a Fast Fourier transform (FFT) on all electrodes. We then separated the signal to the classic frequency bands: delta (1– 3Hz), theta (4 – 7Hz), alpha (8 – 12Hz), beta (13 – 29Hz), and gamma (>30Hz). For each trial (product\commercial viewing), we extracted the average power in each frequency band, in each electrode, resulting in forty different frequency values for each trial (8 electrodes X 5 frequency bands). We used a multiple linear regression to test if the power in the different frequency bands across all electrodes significantly correlated with participants’ WTP values in either Parts 1 or 2, as well as with the *Liking* score measured in Part 2. Furthermore, we used the same model to predict participants’ WTP in part 3 with the neural data from Part 1.

As the number of predictors in each regression was very high (40), we first conducted a dimensionality reduction step using an elastic net. We took only the significant predictors and used them, resulting in a relaxed elastic net regression. To control for multiple comparisons, we applied a Benjamini-Hochberg false discovery rate (FDR) correction to all statistical tests. This procedure was used across all EEG feature analysis reported below.

### Inter-subject correlation

We further examined participants’ ISC scores while they watched products’ pictures (Part 1) and while they watched commercials (Part 2) and tested whether these neural measures from Part 1 were associated to the WTP values stated during the product-viewing session of Part 3. We conducted a cross-correlation analysis to calculate the ISC scores for the different products\commercials. There are several approaches for calculating ISC, and it is used both in fMRI (Hasson et al., 2004) and EEG (Dmochowski et al., 2012; Poulsen et al., 2017). Following Hakim et al. (2021), we computed ISC on time-varying spectral power at 500 Hz. For each participant *i*, stimulus *s* (product-trial in Part 1; commercial in Part 2), frequency band *b* (delta, theta, alpha, beta, gamma), and electrode *e* (8 scalp electrodes), we first formed a leave-one-out group trace by averaging the power time-series of all other participants viewing the same stimulus, at the same band and electrode. We then computed the cross-correlation between participant *i*’s power time-series and the corresponding leave-one-out average across all integer lags and took the peak correlation (maximum over lags) as the electrode-level ISC score for (*i*, *s*, *b*, *e*). This yielded five ISC scores (for five frequency bands) per participant, per stimulus, and per electrode. Finally, to obtain a single ISC value per band, we averaged across electrodes, producing five band-specific ISC scores per participant per stimulus (i.e., per participant×product×trial in Part 1; per participant×commercial in Part 2).

We related valuation measures to the five band-specific ISC scores using linear mixed-effects models, with identical fixed effects but random-effects structures matched to each part’s design. In Part 1, participants viewed and rated each product’s picture twice. i.e., they had two trials per product. We modeled trial-level outcomes (WTP) as *WTP*∼*ISCδ* + *ISCθ* + *ISCα* + *ISCβ* + *ISCγ* with a random intercept for participant×product to account for the two repeated trials within the same participant-product pair. In Part 2, participants viewed each commercial once, with 3-5 commercials per product. We fit the same fixed-effects model but used crossed random intercepts capturing baseline differences among participants, products, and individual commercials.

Additionally, to test whether neural responses to product images in Part 1 could predict participants’ WTP in Part 3, we used the same ISC features from Part 1 (averaged across the two trials per product using Fisher-z transformation and inverse-z back to *r*) to predict Part 3 WTP scores (also averaged across the two trials per product). The model structure remained the same (*WTP*∼*ISCδ* + *ISCθ* + *ISCα* + *ISCβ* + *ISCγ*), with crossed random intercepts for participant and product.

All models were fit by maximum likelihood (L-BFGS). We report RMSE and pseudo-*R*^2^ (marginal *R*^2^*_m_*, variance explained by fixed effects (Nakagawa & Schielzeth, 2013)).

### Frontal Asymmetry

We quantified frontal hemispheric asymmetry (FA) in each frequency band as the log-power difference between the most fronto-lateral electrodes in our setup: *FA* = *ln*(*F*8) − *ln*(*F*7). Band-limited power was computed at every time sample of the power time-series (500 Hz), and FA was evaluated point-wise and then averaged over time, yielding one FA score per participant × band × stimulus (i.e., per participant×product×trial in Part 1; per participant×commercial in Part 2).

To test if FA can predict valuation in Part 1, we fitted a linear mixed-effects model with WTP as the dependent variable and the five band-specific FA scores (delta, theta, alpha, beta, gamma) entered simultaneously as fixed effects. To accommodate the two repeated trials per product, we included a random intercept for each participant×product pair. For Part 2, we used the same fixed-effects specification (five FA bands entered jointly) but a random-effects structure matched to the design: crossed random intercepts for participant, product, and commercial, i.e., (1|*participant*) + (1|*product*) + (1|*commercial*). This was done separately for both WTP and *Liking*.

In addition, we tested whether FA scores from Part 1 were associated with participants’ WTP ratings for the same products in Part 3. To do so, we first averaged the two FA trials per product in Part 1, yielding a single FA score per participant × product × band. We also averaged the two WTP responses per product in Part 3, resulting in one outcome value per participant × product. We then used the same fixed-effects specification (five FA bands entered jointly) to model Part 3 WTP ratings, with a random intercept for each participant × product pair.

### All Features Combined

We also constructed three combined models to evaluate how well EEG features predict valuation across different stages of the experiment. For Part 1 and Part 3 WTP models, we used the EEG features extracted from Part 1 (spectral band power, ISC, and frontal asymmetry), whereas the Part 2 WTP and Liking models used the EEG features extracted from Part 2. For each model, we began with the complete feature set generated for that specific analysis. Following the procedure outlined for the frequency-based models, we first applied an elastic net regression as an initial dimensionality reduction step. All features selected as non-zero predictors by the elastic net were then entered into a multiple linear regression model with standard errors clustered by participant, implementing a relaxed-elastic-net approach in which feature selection and coefficient estimation are performed in separate stages.

### Neural Networks

We implemented a range of neural network models to predict participants’ WTP and *Liking* responses based on EEG data. These include an architecture developed in our lab (DeePay, (Hakim et al., 2023)) alongside established models from prior research: two well-known and frequently used deep learning models, DeepConvNet (Schirrmeister et al., 2017) and EEGNet (Lawhern et al., 2016), as well as state of the art models which are specifically tailored for prediction on EEG data (although not specifically designed for neuromarketing) – EEG_TCNet (Ingolfsson et al., 2020), and TCNet-Fusion (Musallam et al., 2021). To estimate chance-level performance, we included a shuffled-label baseline version of our in-house model architecture (DeePay_TCN_Shuffled).

### Training and testing procedure

We first applied the models to predict WTP in Part 1, replicating a prior study from our lab (Hakim et al., 2023). We then extended this framework to predict both WTP and *Liking* in Part 2, and to predict the WTP of Part 3 based on the EEG data recorded in Part 1. All models were evaluated across multiple prediction conditions, value separation levels, and leave-out procedures (see below).

### Prediction Tasks and Data Splits

We tested the networks’ ability to classify between high and low values (either WTP or *Liking*). See figure 5 for an illustration of the prediction procedure detailed below. In all prediction attempts, we divided the data into 70% training, 15% validation, and 15% test sets. The validation set was used for the hyperparameter tuning procedure and for early stopping during training. As we did in our previous study (Hakim et al., 2023), for training the network in all prediction attempts, we took the bottom 0−35% of values and labeled them as low values, while the top 65−100% was labeled as high values. The rest of the samples in the training set were discarded (35−65%). This was done to boost the network’s ability to identify distinct value information while learning, since the 35−65% of values are the closest in value terms, and hence, would be hardest to distinguish.

**Figure 5.**
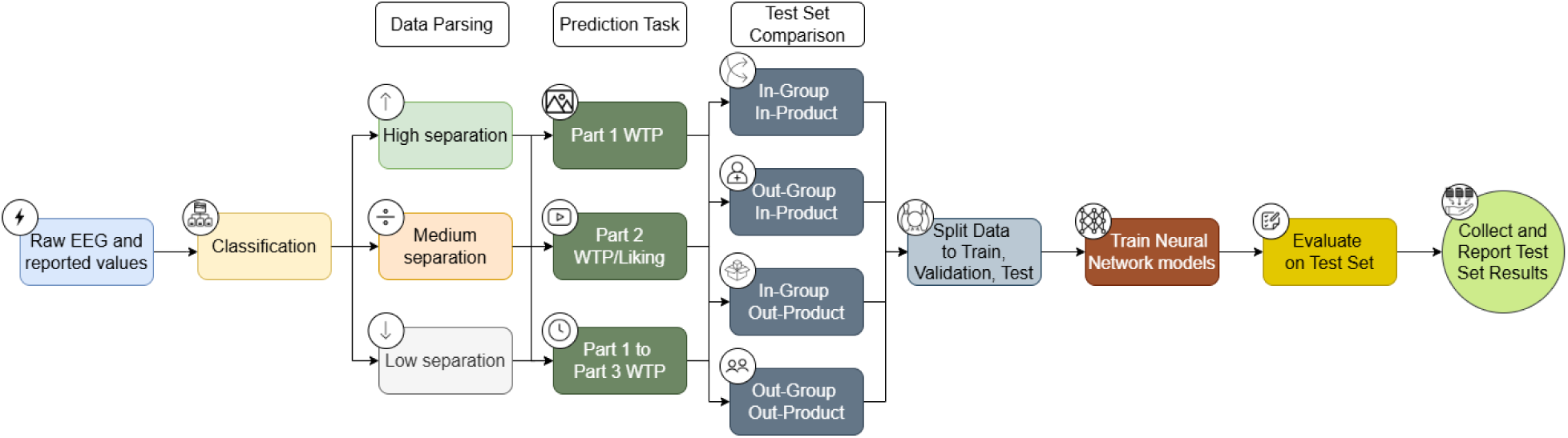
Illustration of the prediction procedure using neural networks.

Based on the prediction condition, we divided the test set such that the high and low values represented a different percentile of the data. We had three different parsing of the data – *High*, *Medium*, and *Low* separations. In the *High* separation, the top 20% of the WTP or *Liking* values and the bottom 20% of values, were the high and low labels for classification, respectively. All other data was not included in the prediction attempt of the test set. In the *Medium* separation, values between the 65%-80% percentiles served as the high labels, and values between 20%-35% percentiles served as the low labels. All the rest of the data was omitted from the test set. In the *Low* separation, values between the 50%-65% percentiles served as the high labels, and values between 35%-50% percentiles served as the low labels of the test set. Again, we omitted all other data from the test set for this prediction.

In all prediction attempts, we report the results of the prediction success of the test set. Following previous work in our lab (Hakim et al., 2023), we expected that the *High* separation will yield the best prediction results. The utility of performing such separations lies in isolating trials that reflect the most extreme and therefore most distinguishable neural valuation states, thereby maximizing the signal-to-noise ratio in the neural data. This approach allows us to test whether the neural network captures robust neural signatures associated with clear preference differences, and to assess how prediction performance generalizes as the distinction between high and low values becomes increasingly subtle.

### Prediction Scenarios Across Parts 1–3

We conducted various predictions on the data. All analyses listed below were conducted using the same

*High, Medium,* and *Low* value separations mentioned above.

First, in Part 1, we attempted to predict participants’ WTP for the products they just saw based on their EEG activity measured while they watched the product images (Part 1 WTP). Second, using the EEG activity measured while participants watched the commercials in Part 2, we attempted to predict participants’ WTP or *Liking* reported right after viewing the commercial (Part 2 WTP/*Liking*). Third, we used the EEG activity measured in Part 1 to predict the WTP for the products participants reported a week later in Part 3 (Part 1 to Part 3 WTP).

### Out-of-Sample/Product Generalization Procedures

We also utilized several different leave-out procedures. Such out-of-sample predictions are critical because it more closely approximates real-world applications, in which models must generalize to new consumers and new advertising materials rather than rely on patterns of the same participants and stimuli. Out-of-sample evaluation also provides an unbiased estimate of model performance and reduces the risk of overfitting (Shmueli, 2010; Yarkoni & Westfall, 2017). Despite this, many neuromarketing studies still rely primarily on within-sample predictions, which can inflate reported performance and limit applicability to industry settings (Hakim et al., 2018).

To address this limitation, we trained the network on one subset of the data and evaluated its predictive performance exclusively on previously unseen stimuli. Specifically, we tested the models using four complementary leave-out procedures: (1) an “In-group In-product” leave-out, where the test set is randomly held out from the pool of all participants and all products; (2) an “In-group Out-product” leave-out, where a randomly selected subset of products (corresponding to 15% of the total trials) was held out for the test set so that the network did not train on those products; (3) an “Out-group In-product” leave-out, where a randomly selected subset of participants (corresponding to 15% of the total trials) was held out for the test set so that the network did not train on those participants; (4) an “Out-group Out-product” leave-out, where a randomly selected subset of participants and products were held out for the test set so that the network did not train on those subjects and products. For the latter three leave-out procedures, the validation set was randomly drawn from the pool of trials remaining after holding out the test set. Furthermore, for the latter three leave-out procedures we iterated over many possible options of participants and products held out for the test set. Note that in the Out-group Out-product procedure, a substantial portion of the data had to be excluded from both the training and test sets. Specifically, we removed all trials in which the *product* was not designated as a held-out product, but the *participant* was held out, as well as all trials in which the *participant* remained in the training pool, but the *product* was held out. Because only trials that simultaneously belonged to held-out participants *and* held-out products could be used for testing (and only trials from the remaining participants and products could be used for training), the effective dataset for this leave-out condition was considerably smaller than in the other procedures, substantially hindering training and prediction. For a full list of the number of trials used in each leave-out procedure see Table W21.

Prior to training and testing, we used the validation set to tune hyper-parameters using the Parzen Estimator approach (Bergstra et al., 2011). These parameters include kernel sizes, number of filters, dropout probability, batch size, and more. Training was carried out with the “Adam” optimizer as implemented in tensorflow and a learning rate of 0.001. Early stopping of the training was specified with a patience of 60 training epochs and utilized categorical cross-entropy as the loss function.

Following established procedures in the field, training was conducted on the train set and validated on the validation set, with the held-out test set used for reported predictions. For each prediction, an ensemble of 5 networks trained with different random initializations was used and reported values correspond to the average prediction of all networks. For classification success, we report accuracy, area under the curve (AUC), and F1 scores. In all prediction attempts, we report the prediction success of the test set (averaged across random initializations), and for the latter three leave-out procedures we report the average values of the prediction scores over all iterations, and the standard error of these averages. We followed the same procedure using all the neural networks mentioned above.

### Equating the time windows for prediction

As participants in Part 1 viewed each picture for 3.5 seconds while they watched each commercial for 6 seconds, we wanted to equate between the amount of predictive data in the different parts. Therefore, in an additional set of analyses, we took 3.5 seconds segments from the EEG activity in Part 2 (commercial viewing) and attempted to predict with them participants’ Part 2 WTP/*Liking*. As the aim of this analysis was to examine the different time segments of the data, for brevity, we performed it only on the In-Group In-Product procedure.

### Classification with logistic regression

To assess how much predictive benefit deep learning provides, we included a benchmark analysis using logistic regression, a widely used classification approach. Logistic regression models were trained on the EEG frequency features described above under the In-Group In-Product condition, where we expected performance to be strongest. Again, separate models were estimated for Part 1 WTP, Part 2 WTP, Part 2 *Liking*, and delayed valuation (Part 1 EEG predicting Part 3 WTP). All regressions were evaluated across the three levels of value separation.

## RESULTS

### Behavioral results

#### BDM procedure

First, we examined if our BDM procedure was successful, and participants demonstrated clear preference ordering of the products so we could use the BDM values in our analyses. Therefore, we examined the overall distribution of the normalized bids across participants in each of the experimental parts. As can be seen in Figure 6, there is a wide and relatively even spread of the WTP values across subjects. This means that most subjects used a wide range of amounts in their bids and did not use only several fixed amounts such as 5, 10 or 20. This shows that our BDM procedure was successful, and participants demonstrated clear preference ordering that we could use in our analyses.

**Figure 6:**
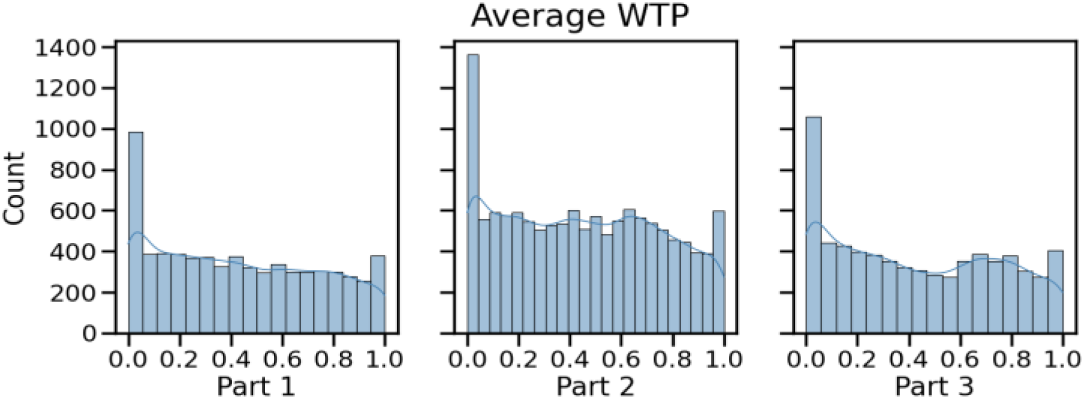
Distribution of average WTP across subjects in each part of the experiment, normalized. The smooth line is an estimated fit of the distribution.

### WTP consistency

Next, we examined the consistency of participants’ bids in each part of the experiment across product repetitions to make sure that the WTP values are reliable and stable values to use for prediction.

Therefore, we calculated the correlation between the first and second bids for each product. As can be seen in Figure 7, we found a strong and significant correlation both in Part 1 (r = 0.8) and in Part 3 (r=0.86). In Part 2, the averaged intraclass correlation coefficient was 0.8. This means that participants were consistent in their bids for a given product across repetitions. Hence, we can consider their average WTP for a given product in a specific part as a valid estimate of their preference for a product.

**Figure 7:**
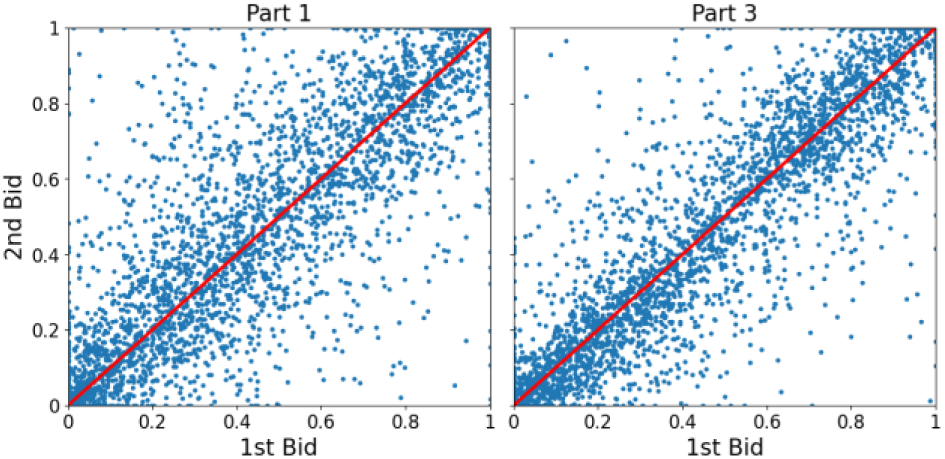
Consistency of WTP values across repetitions of bids. Participants’ WTP for each product across the two repetitions in Part 1 (left) and Part 3 (right). Each blue dot represents the WTP of a product for a given participant in the two repetitions. The red line represents a perfect correlation on the 45° line.

### Commercials’ influence

We fit a linear mixed-effects model to examine whether WTP values changed across the three experimental parts (Part 1, Part 2, Part 3). This design allowed us to test whether exposure to commercials increased WTP immediately (Part 2) and whether such an effect persisted one week later (Part 3). The model included Part as a fixed effect, with random intercepts for participants and items.

We found a significant effect of Part on WTP (see figure 8). Relative to Part 1, WTP was significantly higher in Part 2 (β = 0.018, SE = 0.002, z = 8.62, p < .001), whereas the difference between Part 3 and Part 1 was not significant (β = 0.002, SE = 0.002, z = 1.02, p = .307). Planned pairwise comparisons confirmed that WTP in Part 2 was also significantly higher than in Part 3 (β ≈ 0.016, p < .001, Holm-corrected). These findings indicate that commercials increased participants’ WTP for the advertised products immediately after viewing, but that this effect did not last: one week later, WTP values returned to baseline.

**Figure 8:**
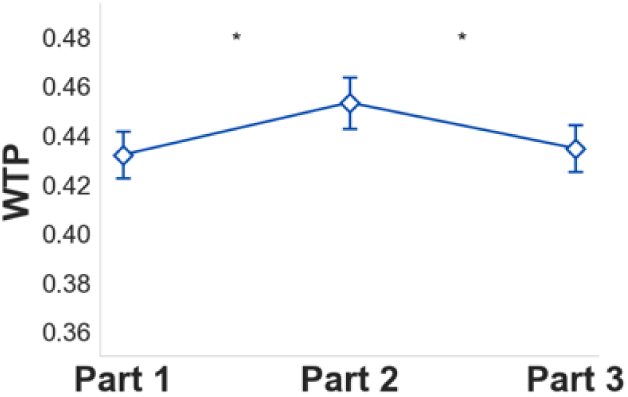
Average WTP (normalized) across all subjects and products in each of the parts.

### Relation between WTP and Liking

We next examined if liking the commercial is significantly correlated with the WTP for the product advertised in the commercial. Therefore, we conducted a simple linear regression on the data from Part 2 where the *Liking* values served as the independent variable and WTP values served as the dependent variable with clustered errors per subject. As can be seen in Table 1, the overall regression was statistically significant (F (1, 110) = 104.4, p < 0.001). Although the relationship was positive and significant, the effect size is relatively small (R^2^ = 0.098). This indicates that enjoying a commercial does not always go hand in hand with perceiving the product featured in it as highly valuable.

**Table 1.** Results of a linear regression for the data in Part 2 between a commercial’s *Liking* value (independent) and the WTP for the product advertised in the commercial (dependent).

|  | Coefficient | Standard error | Z | P-value |
| --- | --- | --- | --- | --- |
| Constant | 0.2879 | 0.018 | 15.740 | 0.000 |
| Liking | 0.3173 | 0.031 | 10.217 | 0.000 |

### EEG Results

Before trying to predict participants’ WTP and *Liking* values using raw EEG as input to a neural network, we used well-known features as predictors in a standard regression. This step served two purposes: it provided an interpretable benchmark grounded in prior literature and ensured that any later improvements from raw EEG deep learning models could be evaluated against a known baseline.

### EEG spectral power

Previous research has demonstrated that specific frequency bands, including alpha, beta, and gamma, can be utilized to predict various marketing metrics, such as a consumer’s liking of a movie or their willingness to pay for a product (Boksem et al., 2025; Boksem & Smidts, 2015; Braeutigam et al., 2004; Hakim et al., 2021; Khushaba et al., 2012, 2013; Koelstra et al., 2012; Luck, 2014; Ravaja et al., 2013; Smith & Gevins, 2004; Telpaz et al., 2015; Xu & Liu, 2024; Yadava et al., 2017). Therefore, we conducted a similar analysis and examined if there is a significant relationship between these traditional predefined EEG frequency bands and WTP values reported in either Parts 1, 2 or 3. To do so, we conducted a multiple linear regression where the average power of the classic frequency bands (of each commercial) served as predictors and WTP values served as the dependent variable (see methods for details). In both Parts 1 and 2, the overall regression was statistically significant but with low R^2^ (Part 1: Adj.R^2^ = 0.009, F30, 129 = 2.43, p < 0.001; Part 2: Adj.R^2^ = 0.013, F37, 123 = 3.826, p < 0.001). In Part 1, we found gamma to be significant in the Fp2, Fz and Pz electrodes. Theta, however, was only significant in Fpz. Lastly, we found delta and beta to be significant in F7. See table W2 for the full regression results. Part 2 showed a slightly different pattern with a significant gamma in Fp1 and Fz. Here, theta was significant in both Cz and Pz, while delta was significant in Fp1, and beta in F7. Unlike Part 1 – here also alpha was significant in F7. See table W3 for the full results.

We further examined the relationship between the traditional EEG frequency bands and commercials’ *Liking* values as measured in Part 2. Here also, the overall regression was statistically significant but with a very low R^2^ (Adj.R^2^ = 0.01, F29, 131 = 2.554, p < 0.001). Here, again, the pattern was not consistent with the other parts. Gamma was significant in Cz, delta in Fz and Cz, and alpha in F8. No other significant predictors were found. See table W4 for the full results.

We next examine the WTP prediction for a week later. To that aim, we tested if the different frequency bands measured in Part 1 are correlated with the WTP for the same products measured a week later in Part 3. Since each product repeats twice, both in Part 1 and in Part 3, and we could not assume the first repetition in Part 1 is necessarily the one to represent the first repetition in Part 3, we averaged the brain activity in Part 1 across repetitions of a given product and also averaged the WTP values across repetitions of that same product appearing in Part 3. Thereafter, we used the relaxed elastic net model for dimensionality reduction leaving eleven predictors. The fitted multiple linear regression model was *WTP_BDM_*_3_ = *β*_0_ + *β*_1_ ∗ *X_BDM_*_1_ + *ε_i_*, *_j_*. The overall regression was statistically significant (Adj.R^2^ = 0.007, F11, 149 = 2.2, p = 0.016), but again with very low R^2^. Here, only three predictors were significant – alpha and delta in F7 and theta in Pz. See table W5 for the full results.

Note, that although significant, the effects sizes in all regressions are very small (∼1% or less explained variance) and the significant predictors and electrodes are not consistent across models precluding any systematic conclusions. This means that although significant, there is only a weak relation between the classic EEG frequency bands and WTP\*Liking* values. Therefore, this calls for employing more advanced non-linear prediction models and to move away from just using the predefined classic frequency bands.

### Inter-subject correlation

Some studies have found that the correlation between participants’ brain activity (inter-subject correlation; ISC) can be used as a measure of engagement towards the stimuli presented (Barnett & Cerf, 2017; H. Y. Chan et al., 2019; Dmochowski et al., 2012, 2014; Hakim et al., 2021; Hasson et al., 2004; Poulsen et al., 2017). Therefore, in both Part 1 and Part 2, we examined the association between band-limited ISC during stimulus viewing and outcome using a linear mixed-effects model. To that aim, we measured the ISC in each frequency band and then used them in one regression model (see methods for specific details of the full procedure). Results of the model fit were for Part 1 *RMSE* = 28.27 with an *R*^2^*_m_* = 0. For Part 2 WTP, the *RMSE* = 27.03 and *R*^2^*_m_* = 0. For *Liking*, *RMSE* = 1.926 and *R*^2^*_m_* = 0.001. See table W6 for the full results of Part 1 and table W7 for the full results of Part 2 – both WTP and *Liking* models. Overall, across all models, none of the predictors were significant except the gamma band ISC for *Liking*. Note also, that the marginal R square is very low in all regressions, calling again for more elaborate methods for prediction. We further tested whether band-limited ISC measured during stimulus viewing in Part 1 predicts WTP assessed one week later in Part 3. The model fit was poor with *RMSE* = 28.37 and *R*^2^*_m_* = 0.003. No ISC band was significant. Full coefficients are reported in table W8. Like Parts 1 and 2, effect sizes were very small, indicating that band-limited ISC provides minimal predictive value for WTP measured a week later.

### Frontal asymmetry

Alpha band asymmetry has been found to be related to approach-avoidance behavior (Cartocci et al., 2016; Davidson, 1998; Koelstra et al., 2012; Ohme et al., 2009, 2010; Ravaja et al., 2013; Sutton & Davidson, 2000; Vecchiato et al., 2010, 2011). Instead of just focusing on alpha asymmetry, and to avoid making any assumptions about the importance of a given frequency band, we calculated the frontal hemispheric asymmetries of all frequency bands, as we previously did (Hakim et al., 2021). Frontal asymmetry was defined per band as ln(*F*8) − ln(F7) and averaged over time within each trial.

In Part 1, we related WTP to the five FA bands using a linear mixed-effects model with a random intercept for each participant×product pair to account for the two repeated trials per product. The results were again significant but poor with *RMSE* = 28.210 and *R*^2^*_m_* = 0.002. Here, alpha, beta, and gamma were significant, though with very low R^2^. The full coefficients are reported in Table W9.

In Part 2, we related WTP (and, in a parallel model, *Liking*) to the five FA bands. Using the same fixed-effects specification, we fit a mixed model with crossed random intercepts for participant, product, and commercial. The results of the model fit for WTP were *RMSE* = 27.01 and *R*^2^*_m_* = 0, and for *Liking RMSE* = 1.922 and *R*^2^*_m_* = 0.003. Here, none of the predictors were significant and the R^2^ was, again, very low. Complete results appear in Table W10.

We next tested whether FA measured during product viewing in Part 1 predicts WTP one week later in Part 3. Again, we averaged FA in Part 1 and WTP in Part 3 across the two repetitions to yield one observation per participant×product, then fit a mixed model with the five FA bands as fixed effects and crossed random intercepts for participant and product. The model fit was again poor with an *RMSE* = 28.304 and *R*^2^*_m_* = 0.007. Here, only gamma was significant, consistent with the significant gamma from Part 1. Though, again, R^2^ was very low. See table W11 for the full results.

### All Features Combined

Following the feature-specific analyses, we examined whether combining all EEG-derived predictors could improve the prediction of the behavioral outcomes. To this end, we used the outputs from all prior analyses – power in each of the frequency bands in each electrode, ISC values, and FA measures. We fit a multiple linear regression after performing an elastic net regression, just like in the frequencies analysis. This was done as a dimensionality reduction step due to the large number of predictors. The analysis was performed separately on the data from Part 1, Part 2 (where both WTP and *Liking* responses were available), and Part 3.

In Part 1, the model was significant, but with a small portion of the variance explained (Adj.R^2^ = 0.008, F4, 160 = 3.402, p < 0.001). Across all predictors, only theta in FPz was significant. For Part 2, separate models were fit for WTP and *Liking*. For WTP, the linear model explained a small but significant portion of the variance (Adj.R^2^ = 0.023, F22, 160 = 3.512, p < 0.001). Across all the predictors, only beta in Fz was significant. None of the ISC or FA predictors were significant. For *Liking*, the model explained slightly less variance (Adj.R^2^ = 0.02, F23, 136 = 3.304, p < 0.001), with no significant predictors across any of the three EEG-derived features. For the delayed valuation, we repeated the combined-predictor analysis using Part 1 features to predict Part 3 WTP. Again, the model was significant but explained only a very small portion of the variance (Adj.R^2^ = 0.016, F11, 160 = 2.781, p = 0.002). See Tables W12-W15 for the full results.

Taken together, all our attempts to predict behavior using standard EEG measures (band power, ISC, and FA or the combination of them) – within-session and across a one-week delay, explained at most ∼0-2% of the variance and yielded inconsistent or null effects. This indicates that these conventional features convey only limited predictive related information. Therefore, these results strengthened our motivation to use more advanced, nonlinear models that incorporates the raw EEG data as the input for analysis without the need to a-priori decide which are the most important features for prediction.

### Deep learning models

As the five neural networks we used produced very similar prediction results in each of the various prediction attempts, and for clarity and being concise, we report in the figures and in the main text the average accuracy across all five different neural network models and compare this average to the model with shuffled labels serving as the null random model. However, see tables W16-W19 for the full details of all prediction metrics separated by neural network model and prediction attempts.

### Part 1 WTP Prediction

In Part 1, we used the EEG activity recorded during passive product viewing (3.5 seconds), before any motor movement, to predict participants’ WTP as reported after viewing each of the products. As can be seen in figure 9 (and table W16 for a detailed description of all prediction results of Part 1), first, across all four leave-out procedures, classification accuracies declined as the value separation declined. That is, the *High* value separation condition consistently yielded the highest prediction accuracies, followed by the *Medium* value separation, while the *Low* value separation yielded the lowest (but still relatively high) prediction accuracies. Second, classification accuracies were the highest for the In-Group In-Product leave-out procedure (74.3%, 67.5%, 57.5%, for *High*, *Medium*, and *Low* value separations, respectively). We achieved similar high classification accuracies in the In-Group Out-Product leave-out-procedure (74.5%, 66.8%, 57.8%, for *High*, *Medium*, and *Low* value separations, respectively). This was followed by also relatively high accuracies for the Out-Group In-Product leave-out-procedure (68.7%, 61.5%, 54.8%, for *High*, *Medium*, and *Low* value separations, respectively), and similarly for the Out-Group Out-Product leave-out-procedure (69.0%, 62.7%, 56.3%, for *High*, *Medium*, and *Low* value separations, respectively). Third, across all leave-out procedures and value separations, all models were significantly above the shuffled-labels model. Importantly, these prediction values are very similar and are a direct replication of our previous study with a similar design of Part 1 (Hakim et al., 2023).

**Figure 9.**
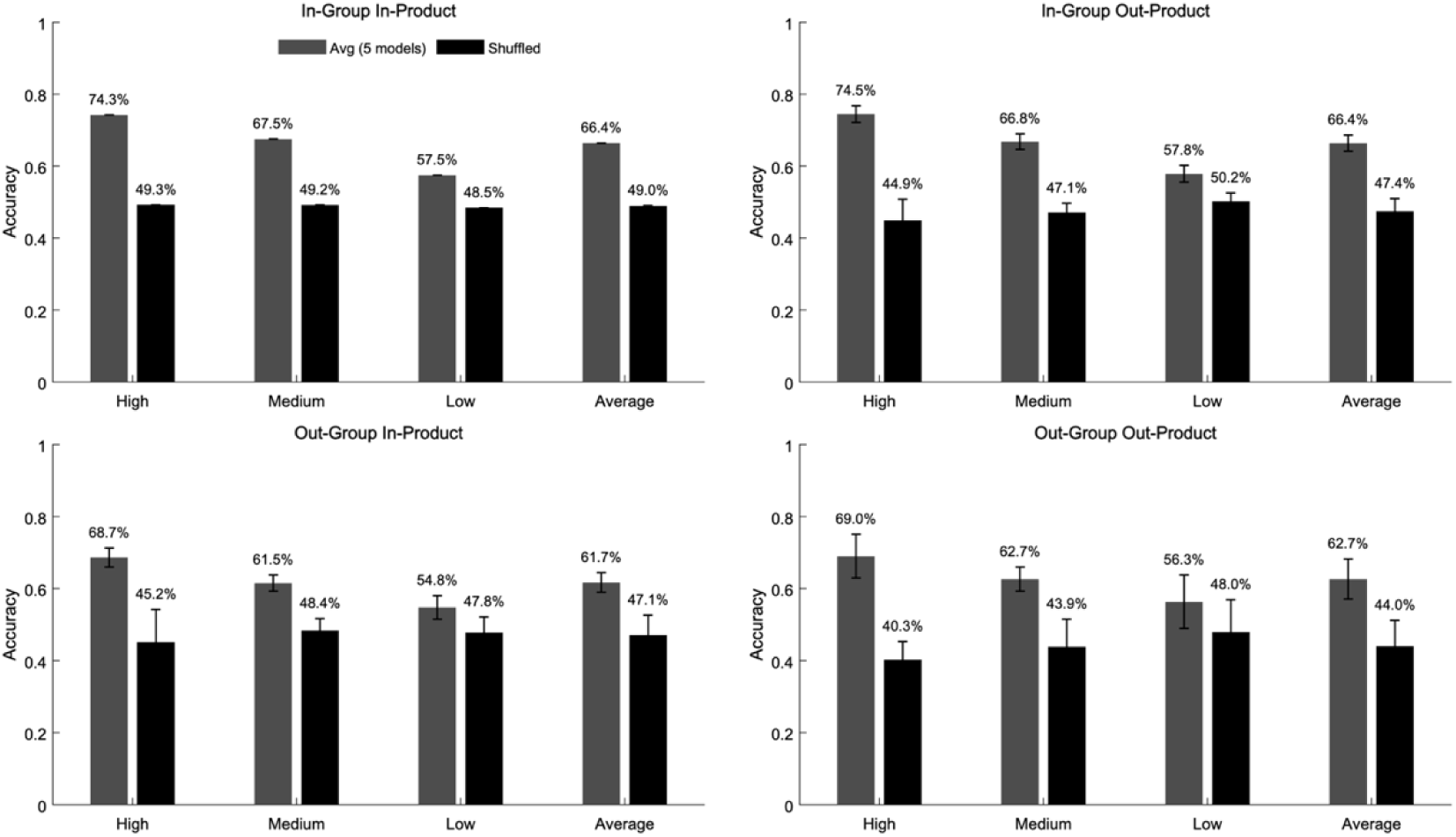
Accuracy results for the WTP prediction in Part 1 in the four leave-out training procedures. A) In-Group In-Product, B) In-Group Out-Product, C) Out-Group In-Product, D) Out-Group Out-Product. The *High*, *Medium* and *Low* value separations are on the X axis along with the Average of the three value separations. Prediction accuracies are the average across the five neural networks used. Error bars represent the standard error of the mean of all model iterations. *Part 2 WTP Prediction*

In Part 2, we used the EEG activity recorded during the 6-second commercial viewing to predict participants’ subsequent reported WTP. As can be seen in figure 10 (and table W17 for a full detailed breakdown of the results), the overall effects of value separations and leave-out procedures were very similar to the effects found in Part 1. However, when comparing each condition, the prediction accuracies were somewhat lower in Part 2 than in Part 1. Specifically, there was between 5%-7% decline in prediction accuracies for the two In-Group leave-out procedures but, importantly, all models were still above the shuffled-labels model (68.8%, 65.1%, 54.8%, for *High*, *Medium*, and *Low* value separations, respectively in the In-Group In-Product and 67.6%, 64.5%, 54.5%, for *High*, *Medium*, and *Low* value separations, respectively in the In-Group Out-Product). On the other hand, for the Out-Group leave-out procedures there was a larger decline of approximately 10% in prediction accuracies resulting in that the prediction accuracies in the *Low* value separation conditions were not different than the shuffled-labels model. Note, however, that using F1 as an alternative prediction metric, we found that even in the *Low* value separations the F1 scores were higher than the model using shuffled labels (see table W17 for details).

**Figure 10.**
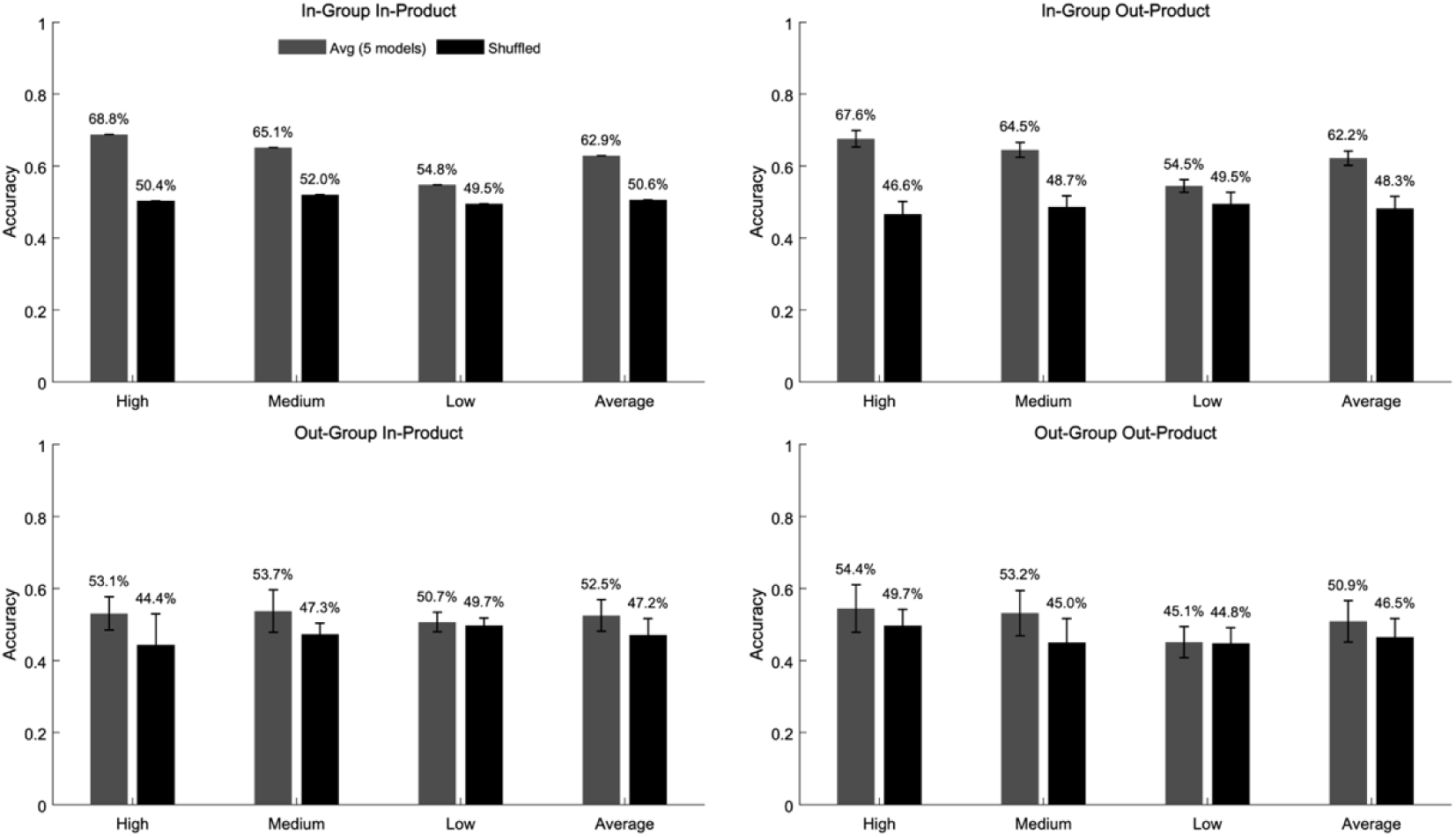
Accuracy results for the WTP prediction in Part 2 across the four leave-out training procedures. A) In-Group In-Product, B) In-Group Out-Product, C) Out-Group In-Product, D) Out-Group Out-Product. The *High*, *Medium* and *Low* value separations are on the X axis along with the Average of the three value separations. Prediction accuracies are the average across the five neural networks used. Error bars represent the standard error of the mean of all model iterations.

Importantly though, the classification accuracies of Part 2 are based on EEG activity while participants watched commercials of the products, while the classification accuracies of Part 1 are based on EEG activity while participants watched images of the products. Although the prediction accuracies were lower compared to Part 1, they were not negligible and significantly higher than the shuffled-labels model. These results underscore the meaningful value-related neural signal captured by the trained models even for a dynamic complex stimulus such as video commercials.

### Part 2 Liking Prediction

We next used the EEG activity recorded during the 6-second commercials viewing to predict participants’ *Liking* responses stated after watching each commercial. As can be seen in figure 11 (see Table W18 for the complete breakdown of prediction results), we were able to successfully predict how much participants liked the commercials in the In-Group leave-out procedures, although, the prediction accuracies were around 5% lower than what we achieved for predicting Part 2 WTP in the same conditions (66.2%, 58.0%, 53.3%, for *High*, *Medium*, and *Low* value separations, respectively in the In-Group In-Product, and 61.4%, 57.2%, 52.4%, for *High*, *Medium*, and *Low* value separations, respectively in the In-Group Out-Product). Here again, we showed the decline in prediction accuracy as the liking separation declines and that all prediction attempts were better than the model with shuffled labels. On the other hand, when moving to the two Out-Group leave out procedures, our prediction attempts were not different than the model with the shuffled labels.

**Figure 11.**
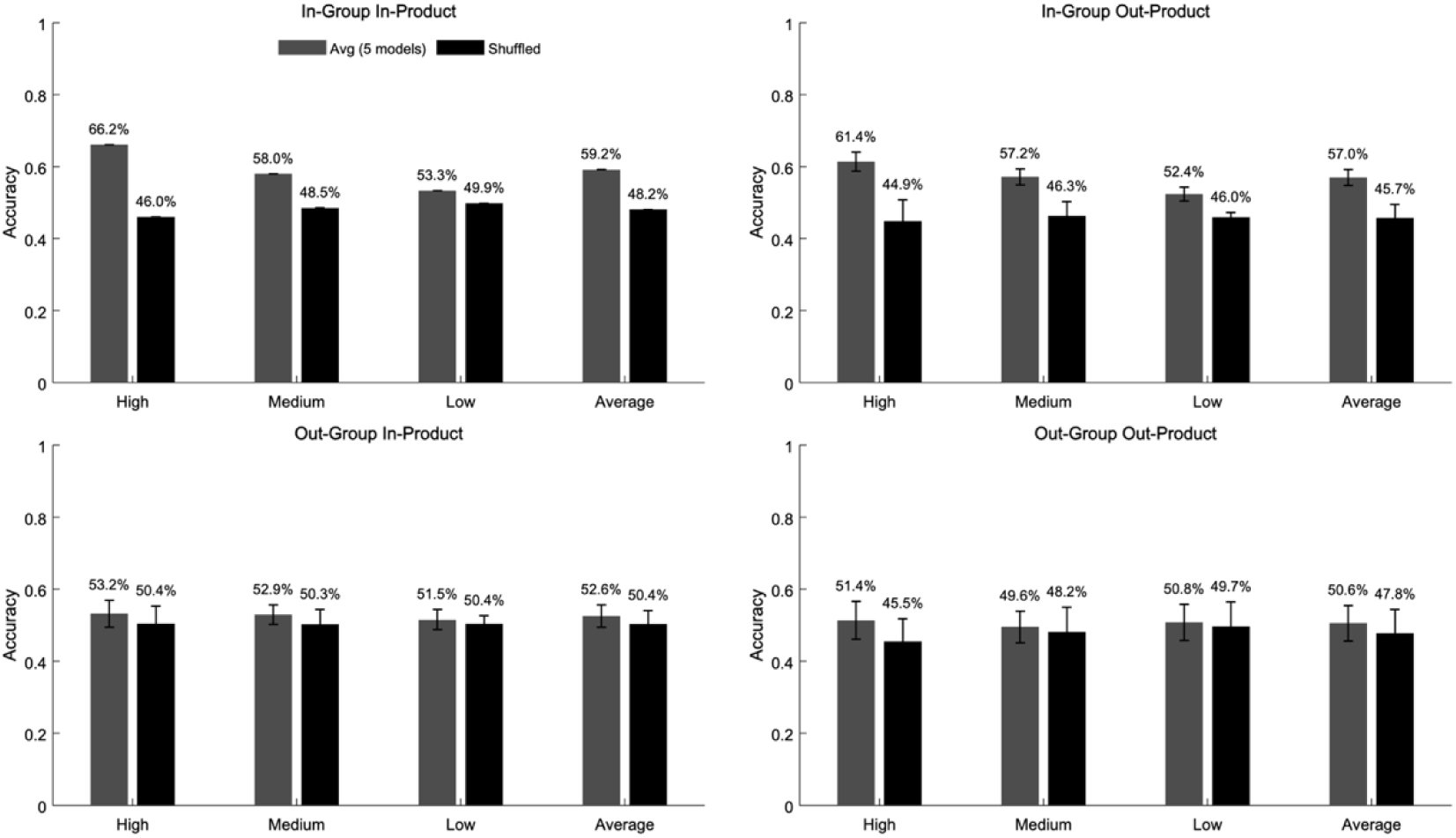
Average accuracy results for the *Liking* prediction in Part 2 across the four leave-out training procedures. A) In-Group In-Product, B) In-Group Out-Product, C) Out-Group In-Product, D) Out-Group Out-Product. The *High*, *Medium* and *Low* value separations are on the X axis along with the Average of the three value separations. Prediction accuracies are the average across the five neural networks used. Error bars represent the standard error of the mean of all model iterations.

### Part 1 EEG activity to Part 3 behavior

We next examined whether EEG activity recorded during product viewing in Part 1 could predict participants’ product preferences measured one week later, as reflected in their Part 3 WTP responses. This analysis tested the long-term predictive value of the neural signals and paralleled the structure of Part 1 predictions by evaluating all four leave-out procedures and the three value-separation levels and using the shuffled DeePay_TCN model as the chance-level baseline.

As can be seen in figure 12 (and full details in table W19), the pattern of results largely mirrored the findings of Part 1 and Part 2. As before, the *High* value separation condition produced the strongest predictions, followed by *Medium*, and then *Low* value separations. Again, prediction accuracies were also dependent on the leave-out procedure. Performance decreased systematically as the generalization demands increased where the In-Group In-Product yielded the highest accuracies, slightly better than the In-Group Out-Product procedure followed by the Out-Group In-Product, and finally the Out-Group Out-Product having the lowest prediction accuracies. However, classification accuracies in Part 3 were consistently lower across all conditions than in the within-session predictions of Part 1, reflecting the increased difficulty of predicting valuation shifts that emerge after a week delay.

**Figure 12.**
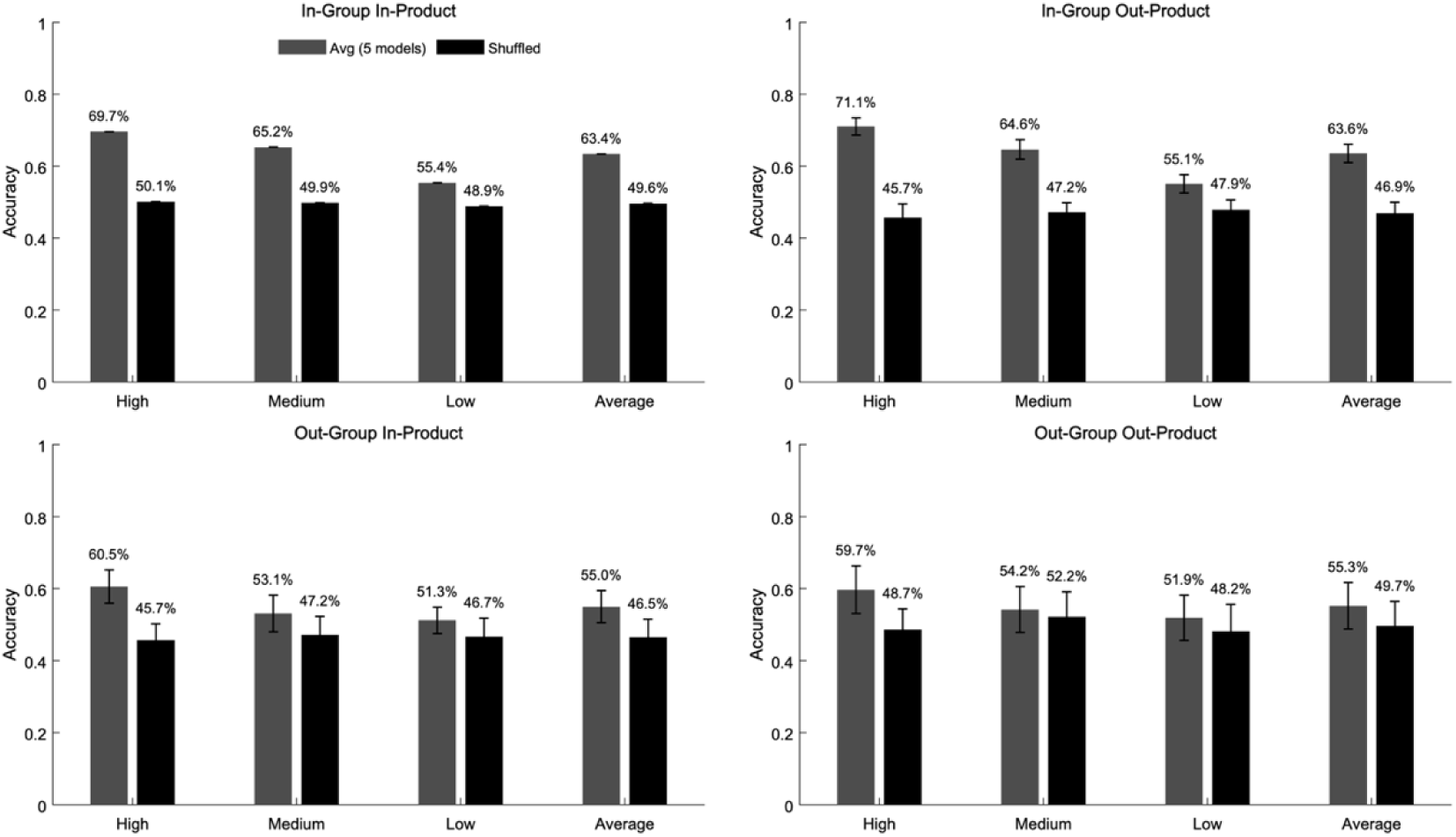
Average accuracy results for the WTP prediction in Part 3 from Part 1 EEG data across the four leave-out training procedures. A) In-Group In-Product, B) In-Group Out-Product, C) Out-Group In-Product, D) Out-Group Out-Product. The *High*, *Medium* and *Low* value separations are on the X axis along with the Average of the three value separations. Prediction accuracies are the average across the five neural networks used. Error bars represent the standard error of the mean of all model iterations.

### Is there a significant part in the data for prediction?

As the time window for analysis in Part 1 and Part 2 was different, we conducted an additional analysis to ensure comparability between the two parts. Therefore, we conducted a sliding window analysis in which we used only 3.5-second segments from the 6-second EEG recordings during commercials viewing in Part 2 yielding six time-window segments. This matched the duration of the product image viewing in Part 1 and allowed us to evaluate whether shorter segments of EEG data during dynamic commercials could yield comparable predictive performance. For brevity, this analysis was performed only on WTP under the *High* value separation and only in the In-Group In-Product condition.

As can be seen in figure 13, performance remained consistent across all time windows. The average classification accuracy for WTP was 67% across the six time-window segments analyzed, which is far better than chance but lower than the prediction accuracies of this condition in Part 1. Critically, no specific 3.5-s window yielded systematically higher accuracy, F1 score, or AUC. That is, we observed no early-, mid-, or late-segment advantage. These results demonstrate that even short segments of EEG activity during commercial viewing contain sufficient information to predict preferences, consistent with those observed for static image viewing in Part 1 (albeit slightly lower).

**Figure 13.**
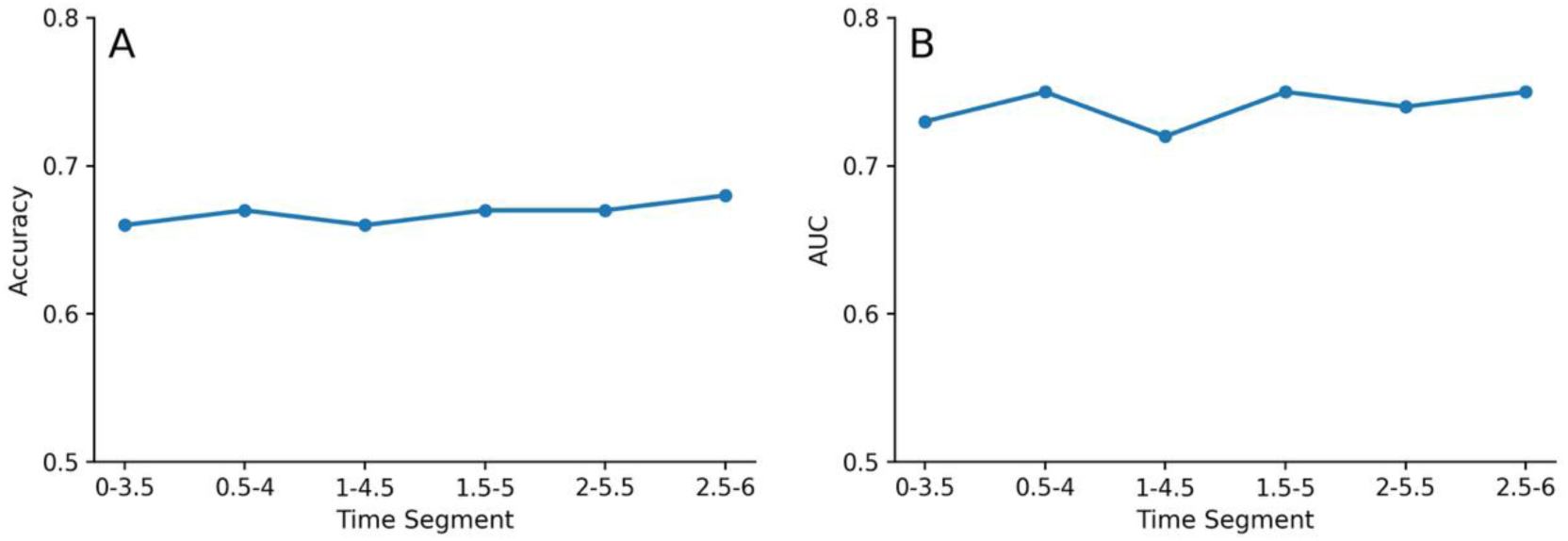
Sliding-window EEG prediction performance of WTP in Part 2. Accuracy (A) and AUC (B) for WTP prediction using six sliding 3.5-s windows extracted from the 6-s duration of the commercials in Part 2. Analysis was conducted on for the *High* value separation, In-Group In-Product. Performance remained stable across all windows, with no early, middle, or late segment showing a systematic advantage.

### Classification with logistic regression

As a benchmark, we compared the prediction performance of the neural network models to logistic-regression classifiers trained separately on frequency-band power, ISC, FA, or their combination.

Again, for brevity, analyses were restricted to the In-Group In-Product condition and evaluated at *High*, *Medium*, and *Low* value-separation levels. We conducted separate models for Part 1 WTP, Part 2 WTP, Part 2 *Liking*, and delayed valuation (Part 1 features predicting Part 3 WTP). We report test accuracy, with the majority-class (chance) accuracy in parentheses; binomial p-values test accuracy above chance, and AUC permutation p-values test AUC above 0.5.

As can be seen in figure 14 (and details in table W20), across feature sets and experimental parts, logistic-regression performance was weak and largely close to chance, with only small and inconsistent above-chance accuracies, almost exclusively at *High* and occasionally *Medium* value separations.

**Figure 14.**
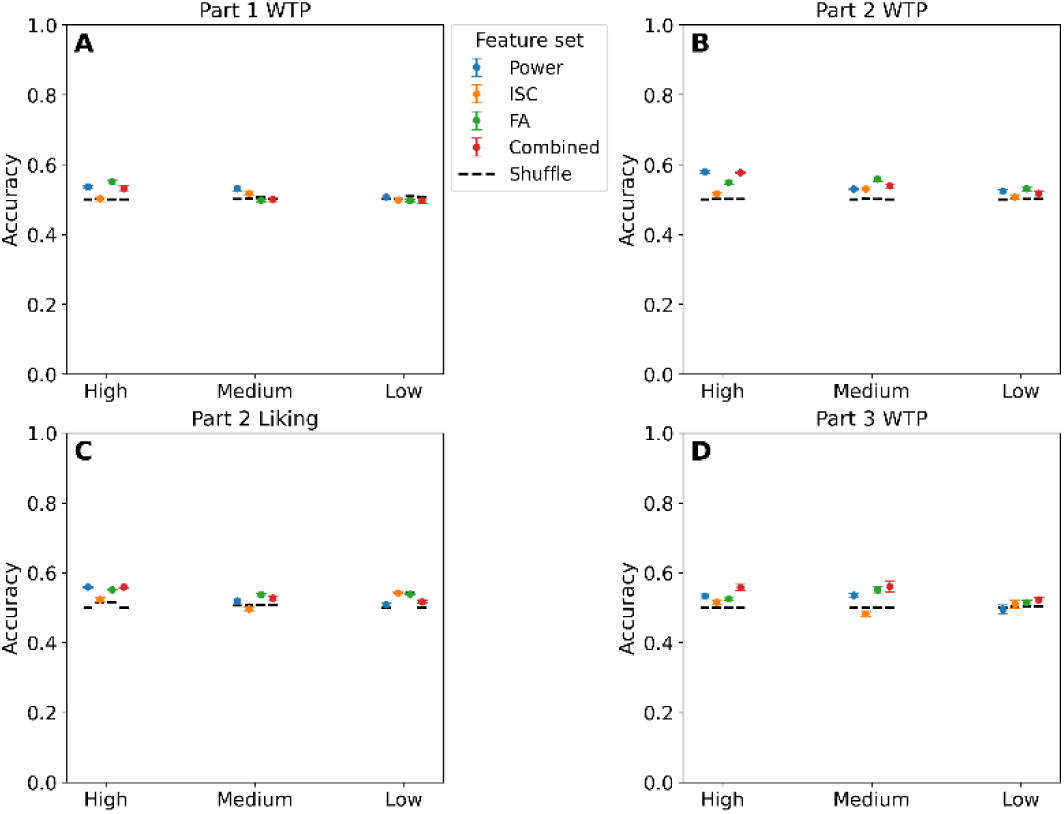
Logistic-regression classification accuracies across EEG feature sets and value-separation levels. Each panel (A– D) shows classification accuracy (mean±SE) for logistic-regression models trained on one of four EEG feature sets – frequency-band power (power), inter-subject correlation (ISC), frontal asymmetry (FA), and their combined predictor set – evaluated at the *High*, *Medium*, and *Low* value separation levels (Combined). Dots indicate mean accuracy and error bars denote standard error. Short black horizontal lines indicate chance level for each condition separately. A) Part 1 WTP, B) Part 2 WTP, C) Part 2 *Liking*, and D) delayed valuation (Part 1 EEG predicting Part 3 WTP). Across conditions, accuracies remain close to 50%, with Power and FA producing the strongest (though modest, ∼57%) above-chance performance in the *High* value separation for most leave-out procedures and conditions. ISC and Combined features generally perform similarly or slightly below Power and FA. As value separation decreases from *High* to *Low*, accuracies for all feature sets trend downward or flatten toward chance, indicating that models using standard EEG features capture only weak and inconsistent decision-related signals. These results provide a baseline comparison for the substantially higher performance achieved by the neural-network models.

Frequency-band power produced the most reliable improvements – e.g., Part 2 WTP reached 57.2% (*High*) and 55.8% (*Medium*) compared to a 50% chance level, and delayed valuation achieved 53.7-54.4% at *High*/*Medium* separations. FA showed similar but slightly weaker effects, with modest above-chance performance in Part 1 WTP and Part 2 WTP at *High* separation. ISC was the weakest feature type, remaining near chance across most tasks except for a small boost in Part 2 Liking at *High* value separation. Combined-feature models improved performance somewhat in Part 2 WTP and Part 2 *Liking* at *High*/*Medium* value separation but again fell to chance at *Low* value separation.

Critically, all logistic-regression models converged to chance as separation narrowed, with *Low* value separation accuracies almost universally between 49-52%, regardless of feature set or part. By contrast, the neural-network models achieved substantially higher and more reliable classification performance across all parts and separation levels. This striking contrast supports the view that value-related EEG signatures are distributed, high-dimensional, and non-linear-properties that are poorly captured by linear classifiers using a-priori set of EEG features but are effectively leveraged by deep learning models without using any predefined EEG features.

## DISCUSSION

In this study, using a large data set of 161 subjects, each viewing multiple different products (n=26) and commercials for those products (n=100), we demonstrate that EEG signals recorded during both the initial product exposure (Part 1) and subsequent commercial viewing (Part 2) can be used to predict two consumer-relevant outcomes at the individual level. Neural activity during Part 1 predicted participants’ initial product valuations, while neural activity during Part 2 predicted how much they would be willing to pay for the product after seeing a commercial advertising the same product they saw in Part 1. Remarkably, brain responses during the first product exposure also anticipated how much participants valued the product a week later, suggesting that early impressions leave a measurable neural trace with lasting value-related predictive information. In addition, our models were able to predict how much participants liked the commercials themselves. Together, these findings show that EEG recordings from both product and commercial viewing contain meaningful signals that can anticipate how consumer’s preferences will evolve, both immediately and over time.

The deep neural network models we used showed robust but variable accuracy in predicting different consumer response outcomes. Performance depended on the out-of-sample generalization procedure, the differences in value between the predicted stimuli, the target variable for prediction, and the time since EEG measurement.

Overall, the prediction results demonstrate that both value separation and leave-out procedure affect prediction performance. As expected, performance was strongest (∼70%-75% accuracy) when the training and test data shared both subjects and products (In-Group In-Product). Although this is considered an out-of-sample prediction due to the use of train and test sets, this leave-out procedure is relatively the easiest. This is because the model is trained on EEG data from all participants and all products allowing it to identify unique EEG characteristics of all products and participants, which help when attempting to predict items from the test set that share these same unique characteristics.

Nevertheless, we managed to show meaningful out-of-sample generalization. The deep learning models generalized well and showed similar high classification accuracies (also ∼70%) to new products when trained on the same participants (In-Group Out-Product). Moreover, classification accuracies on unseen participants when trained on all products (Out-Group In-Product) were slightly lower but still high (∼67%). Therefore, we show that the deep learning models can generalize better for new products when trained on all participants compared to when trained on all products but need to generalize for new participants. This suggests that the unique variability and neural characteristics of a participant is larger compared to the neural valuation signature of a product across participants, which decreases the ability of the models to generalize for new participants compared to new products. Not surprisingly, the classification accuracies were the lowest (but still above chance) when the models attempted to generalize for a new product evaluated by a new participant (Out-Group Out-Product). However, note that for this leave-out procedure, the amount of training and mainly test data were substantially lower than all other cases, limiting the possible prediction success (see table W21 for full details).

Importantly, as we have previously demonstrated (Hakim et al., 2023), performance also scaled with value separation distance. Prediction accuracies were highest (up to 75% in the In-Group In-Product) when the value distance between the products was the largest (*High* value separation), intermediate under *Medium* separation, and lowest under *Low* separation. Importantly, however, all value separations were still better than shuffled labels. This monotonic relation indicates that when the behavioral categories are farther apart in value space, the underlying neural representations are more discriminable, yielding stronger predictions. As value distance shrinks, neural patterns increasingly overlap and classification accuracies approaches but remains above shuffled labels. This suggests that EEG models are particularly good at identifying consumers who will strongly value or devalue a product but are less effective at picking up subtle or moderate differences. A possible approach to increase prediction accuracies when comparing between similar valued products/commercials is to present the same stimuli many times and therefore, reduce noise and increase signal to noise ratio, which will increase prediction accuracies.

In addition, the best performance was for predicting WTP for products when the EEG measurements were acquired while participants viewed pictures of the products (Part 1) compared to viewing commercials that advertised those same products (Part 2). This is probably because participants had clearer and less noisy neural value representations when just viewing the relevant product to be predicted compared to viewing a commercial for that same product. Commercials contain dynamic audiovisual streams, narrative progression, emotional cues, and ongoing persuasive messaging, all of which may produce dynamic high-dimensional neural activity patterns that vary across individuals and induce large changes in valuation while watching the commercial. This added complexity makes it harder for models to extract stable value-relevant features from EEG data during commercial exposure. Note, that even when we used shorter time windows from the commercials, comparable to the length of product viewing in Part 1, it did not improve prediction in Part 2.

We also showed that not every value related measurement yields similar results. Specifically, although there was a positive link between how much participants liked a commercial and how much they valued the advertised product in it, the relationship was relatively weak (in sample R^2^ was less than 10%). This indicates that enjoying a commercial does not necessarily mean that the advertised product will be seen as more valuable. Moreover, in Part 2, predicting commercials’ liking was notably harder than predicting products’ WTP. The best accuracy we achieved was ∼68% (In-Group In-Sample, and In-Group Out-Sample), with reduced generalizability to unseen participants. The comparatively lower predictability of liking can be attributed to two aspects of the experimental design. First, during commercial viewing, participants were explicitly instructed to think about how much they value the product advertised in the commercial, which likely amplified valuation-related neural processes for the product at the expense of purely affective appraisal of the entire commercial. Second, because liking and WTP are only modestly associated – i.e., they overlap only partially, constrains how well value-tuned EEG features can recover variance in liking, thereby yielding lower predictability for this outcome. Last, liking a commercial is affected by various and numerous affective, sensory, and cognitive components, which vary across participants and commercials. Hence, the neural network models had a harder time to capture all these components relative to an overall value representation.

In addition to predicting preferences stated a few seconds after stimulus presentations, we also wanted to examine if we could predict preferences stated a week later after stimulus presentation and EEG measurements. On the behavioral level, participants generally valued products more after viewing the commercials. However, this boost in WTP tended to diminish over time, with product valuations largely returning to their initial baseline a week later. Importantly, using the neural activity measured during initial product viewing (Part 1), we managed to successfully predict the WTP for the same products a week later (Part 3). Note, however, that overall, the prediction performance was lower in Part 3 compared to Part 1 but still better than chance across all conditions, demonstrating that neural patterns recorded during brief product viewing continue to contain meaningful predictive information about later valuation. Notably, even the most demanding leave-out settings – requiring generalization to both new participants and new products – yielded, in most cases, accuracies that surpassed chance, indicating that the neural representations of value measured in Part 1 retain discriminative power over extended time scales. Taken together, these findings show that EEG signals recorded during initial exposure to products carry stable, temporally durable signatures that enable forecasting of consumer preferences expressed a week later. To our knowledge, this represents the first evidence that neural responses to advertising can successfully predict individual WTP at such a long temporal delay.

Using well-known and predefined EEG features for prediction, such as frequency band powers, ISC, and FA, yielded significant results in the regression models. However, importantly, the effect sizes (as measured by the explained variance of the regression models) were extremely low (0-2%) across all models and features. Moreover, the features that were significant and their electrode location varied considerably across the regression models, which prevented us from coming up with coherent conclusions regarding the importance of specific EEG features for value prediction. In addition, using standard logistic regression models with these predefined features yielded prediction accuracies that were mostly not different than chance (∼50-54%). This suggests that neural value representations for products and commercials are more complex, dynamic, and non-linear and cannot be captured by using these a-priori EEG features. Hence, for achieving higher and meaningful prediction accuracies we need to use models that can deal with the complexity of the raw EEG data. As we have previously shown (Hakim et al., 2023), and extend here, using deep neural networks is a suitable approach for analysis.

This study offers several methodological strengths. First, our models were evaluated using rigorous out-of-sample and out-of-product predictions, achieving approximately 65%-70% accuracy in forecasting valuation outcomes. This demonstrates that EEG signals contain generalizable information beyond specific participant-product pairings. In practical terms, the approach can be applied to the two cases that matter outside the lab: predicting responses for new individuals and for new products that were not in the training set. This moves our method towards real-world scenarios, where audiences and content change continuously rather than remaining tied to fixed participant-product pairs. Second, our sample was a lot larger (N = 161) than most previous EEG studies in the field. Third, our study included individuals aged 20 to 63 with diverse educational and occupational backgrounds, enhancing external validity compared to many prior studies that rely heavily on student samples.

At the same time, certain methodological choices warrant further investigation. Participants were instructed to consider their WTP while viewing the product images and commercials, which likely emphasized valuation-related neural processes at the expense of more affective responses. This framing may explain why our models predicted WTP more accurately than commercial liking. Future work could compare different instructional framings to assess their influence on predictive signal quality.

Additionally, all stimuli were presented in a controlled laboratory environment, minimizing distractions and enhancing internal validity. However, this setting may limit ecological realism. To assess the practical utility of EEG-based prediction tools, future studies should evaluate whether similar performance can be achieved in more naturalistic contexts such as ad viewing at home, on mobile devices, or in everyday multitasking environments.

Furthermore, although we used deep learning models optimized for EEG data that demonstrated strong predictive accuracy, their interpretability remains limited. However, our attempts to predict valuation using conventional interpretable EEG features such as frontal asymmetry, ISC, or classic spectral power, yielded poor results. This suggests that valuation-related information is encoded in complex, nonlinear patterns that standard metrics fail to capture. As such, deep learning models appear to be both advantageous and essential for unlocking the predictive potential of EEG in this domain. Future work should continue exploring interpretability tools that can reveal which aspects of the neural signals drive predictions, enabling both theoretical insights and more transparent applications.

Finally, while our models predict valuation, they do not explain why a given participant experiences a shift in valuation as a function of commercial viewing or time. The direction and magnitude of persuasion likely depend on a range of factors, such as emotional resonance, memory encoding, personal relevance, and external factors like the commercial being watched. These factors may be partially accessible through complementary measures like eye tracking or facial expression analysis. Future research should explore multimodal approaches that combine EEG with other behavioral and physiological signals to improve both accuracy and insight (Perez et al., 2024).

Our findings advance the broader effort to understand how advertising reshapes consumers’ preferences and subsequent behavior, and how those shifts can be detected at the neural level. While much previous work has focused on group-level averages or post-hoc correlations between brain responses and outcomes, we demonstrate that individual participants’ EEG data, collected while they viewed a product or a commercial, can be used to anticipate their future preferences. Importantly, this includes not only immediate valuation shifts but also changes that endure for at least a week – a time window relevant for real-world consumer decision-making. This temporal dimension, combined with the individual-level resolution, sets the present approach apart from traditional techniques that often rely on stated intentions or retrospective reports.

Moreover, the fact that predictive information was present in EEG responses during both product and commercial viewing phases suggests that consumer valuation is constructed through multiple cognitive stages. Neural responses to the product before any commercial exposure already predict WTP, highlighting the role of early impressions. Separately, EEG during commercial exposure also predicted valuation indicating that persuasive effects are not confined to the moment of product viewing or ad exposure alone. This supports a view of persuasion as a temporally distributed process, with both initial perceptions and subsequent advertising experiences shaping consumer value.

From an applied perspective, these findings open the door to practical tools for testing and optimizing commercials. EEG-based prediction can help identify which product–commercial combinations are likely to generate meaningful shifts in value for specific individuals. Notably, we achieved these predictions using an 8-electrode system – a configuration aligned with many portable, industry-friendly EEG solutions. This suggests that neuroscience-informed ad testing is not only feasible but scalable, making it attractive for real-world deployment in pre-market evaluations, concept screening, and targeted campaign development.

Beyond methodological innovation, these insights also have practical and ethical implications for the marketing world. Understanding how and when commercials leave lasting neural traces can inform smarter decisions about ad frequency and timing. For instance, whether repeated exposures are necessary to build brand loyalty or whether a single impression is sufficient. Better predictive tools across different content types (e.g., static product images vs. full commercials) can also help marketers allocate budgets more efficiently, ensuring consumers encounter ads that feel relevant and meaningful. Importantly, predictive neuromarketing tools need not serve only commercial efficiency, they can also support more ethical practices, allowing companies to design ads that empower rather than exploit, offering genuine value and fostering trust-based relationships with their audiences.

Together, these findings show that non-invasive EEG signals combined with state-of-the-art deep learning models can be used to predict meaningful, individual-level changes in product valuation induced by advertising, not only immediately, but also after a delay of one week. By combining neurophysiological data with deep learning models, we offer a temporally sensitive and behaviorally relevant view of marketing that complements both traditional marketing measures and prior neuroimaging research. More broadly, this work contributes to a growing literature that seeks to understand and anticipate how consumer preferences evolve over time. As neuroscience continues to intersect with marketing, tools like EEG may play an increasingly central role in revealing the hidden processes that shape consumer decisions.

## Supporting information

full supplemetary file

## References

1. Agarwal, S., & Dutta, T. (2015). Neuromarketing and consumer neuroscience: Current understanding and the way forward. DECISION 2015 42:4, 42(4), 457–462. 10.1007/S40622-015-0113-1

2. Aldayel, M., Ykhlef, M., & Al-Nafjan, A. (2020). Deep Learning for EEG-Based Preference Classification in Neuromarketing. Applied Sciences 2020, Vol. 10, Page 1525, 10(4), 1525. 10.3390/APP10041525

3. Baldo, D., Parikh, H., Piu, Y., & Müller, K. M. (2015). Brain Waves Predict Success of New Fashion Products: A Practical Application for the Footwear Retailing Industry. 10.1177/2394964315569625, 1(1), 61–71. https://doi.org/10.1177/2394964315569625

4. Barnett, S. B., & Cerf, M. (2017). A Ticket for Your Thoughts: Method for Predicting Content Recall and Sales Using Neural Similarity of Moviegoers. Journal of Consumer Research, 44(1), 160–181. 10.1093/JCR/UCW083

5. Becker, G. M., DeGroot, M. H., & Marschak, J. (1964). Measuring utility by a single-response sequential method. Behavioral Science, 9(3), 226–232. 10.1002/BS.3830090304

6. Bergstra, J. S., Bardenet, R., Bengio, Y., & Kégl, B. (2011). Algorithms for Hyper-Parameter Optimization.

7. Berns, G. S., & Moore, S. E. (2012). A neural predictor of cultural popularity. Journal of Consumer Psychology, 22(1), 154–160. 10.1016/J.JCPS.2011.05.001

8. Bilucaglia, M., Mainardi, L., Ramsøy, T. Z., Zak, P. J., Zito, M., & Russo, V. (2025). Editorial: Machine-learning/deep-learning methods in neuromarketing and consumer neuroscience. Frontiers in Human Neuroscience, 19, 1638225–1638225. 10.3389/FNHUM.2025.1638225/BIBTEX

9. Boksem, M. A. S., & Smidts, A. (2015). Brain Responses to Movie Trailers Predict Individual Preferences for Movies and Their Population-Wide Commercial Success. Journal of Marketing Research, 52(4), 482– 492. 10.1509/jmr.13.0572

10. Boksem, M. A. S., van Diepen, R. M., Eijlers, E., Boekel, W., & Smidts, A. (2025). Do EEG Metrics Derived from Trailers Predict the Commercial Success of Movies? A Systematic Analysis of Five Independent Datasets. Journal of Marketing Research, 62(4), 703–720. 10.1177/00222437241309875

11. Braeutigam, S., Rose, S. P. R., Swithenby, S. J., & Ambler, T. (2004). The distributed neuronal systems supporting choice-making in real-life situations: Differences between men and women when choosing groceries detected using magnetoencephalography. European Journal of Neuroscience, 20(1), 293–302. 10.1111/J.1460-9568.2004.03467.X

12. Byrne, A., Bonfiglio, E., Rigby, C., & Edelstyn, N. (2022). A systematic review of the prediction of consumer preference using EEG measures and machine-learning in neuromarketing research. Brain Informatics 2022 9:1, 9(1), 1–23. 10.1186/S40708-022-00175-3

13. Calvert, G. A., & Brammer, M. J. (2012). Predicting consumer behavior: Using novel mind-reading approaches. IEEE Pulse, 3(3), 38–41. 10.1109/MPUL.2012.2189167

14. Cartocci, G., Cherubino, P., Rossi, D., Modica, E., Maglione, A. G., Di Flumeri, G., & Babiloni, F. (2016). Gender and Age Related Effects while Watching TV Advertisements: An EEG Study. Computational Intelligence and Neuroscience, 2016. 10.1155/2016/3795325

15. Castellion, G., & Markham, S. K. (2013). Perspective: New Product Failure Rates: Influence of Argumentum ad Populum and Self-Interest. Journal of Product Innovation Management, 30(5), 976–979. 10.1111/J.1540-5885.2012.01009.X

16. Chan, H. Y., Smidts, A., Schoots, V. C., Dietvorst, R. C., & Boksem, M. A. S. (2019). Neural similarity at temporal lobe and cerebellum predicts out-of-sample preference and recall for video stimuli. NeuroImage, 197, 391–401. 10.1016/J.NEUROIMAGE.2019.04.076

17. Chan, H.-Y., Boksem, M. A. S., Venkatraman, V., Dietvorst, R. C., Scholz, C., Vo, K., Falk, E. B., & Smidts, A. (2024). Neural Signals of Video Advertisement Liking: Insights into Psychological Processes and Their Temporal Dynamics. Journal of Marketing Research, 61(5), 891–913. 10.1177/00222437231194319

18. Christoforou, C., Papadopoulos, T. C., Constantinidou, F., & Theodorou, M. (2017). Your Brain on the Movies: A Computational Approach for Predicting Box-office Performance from Viewer’s Brain Responses to Movie Trailers. Frontiers in Neuroinformatics, 11(72). 10.3389/FNINF.2017.00072

19. Costa-Feito, A., González-Fernández, A. M., Rodríguez-Santos, C., & Cervantes-Blanco, M. (2023). Electroencephalography in consumer behaviour and marketing: A science mapping approach. Humanities and Social Sciences Communications, 10(1), 1–13. 10.1057/S41599-023-01991-6;SUBJMETA=4000,4001,4014;KWRD=BUSINESS+AND+MANAGEMENT

20. Crawford, C. M. (1977). Marketing Research and the New Product Failure Rate. 10.1177/002224297704100216, 41(2), 51–61. https://doi.org/10.1177/002224297704100216

21. Davidson, R. J. (1998). Anterior electrophysiological asymmetries, emotion, and depression: Conceptual and methodological conundrums. Psychophysiology, 35(5), 607–614. 10.1017/S0048577298000134

22. Delorme, A., & Makeig, S. (2004). EEGLAB: an open source toolbox for analysis of single-trial EEG dynamics including independent component analysis. Journal of Neuroscience Methods, 134(1), 9–21. 10.1016/J.JNEUMETH.2003.10.009

23. Dimpfel, W. (2015). Neuromarketing: Neurocode-Tracking in Combination with Eye-Tracking for Quantitative Objective Assessment of TV Commercials. Journal of Behavioral and Brain Science, 05(04), 137–147. 10.4236/JBBS.2015.54014

24. Dmochowski, J. P., Bezdek, M. A., Abelson, B. P., Johnson, J. S., Schumacher, E. H., & Parra, L. C. (2014). Audience preferences are predicted by temporal reliability of neural processing. Nature Communications 2014 5:1, 5(1), 1–9. 10.1038/ncomms5567

25. Dmochowski, J. P., Sajda, P., Dias, J., & Parra, L. C. (2012). Correlated components of ongoing EEG point to emotionally laden attention—A possible marker of engagement? Frontiers in Human Neuroscience, 6(MAY 2012), 23556. 10.3389/FNHUM.2012.00112/BIBTEX

26. Esch, F. R., Möll, T., Schmitt, B., Elger, C. E., Neuhaus, C., & Weber, B. (2012). Brands on the brain: Do consumers use declarative information or experienced emotions to evaluate brands? Journal of Consumer Psychology, 22(1), 75–85. 10.1016/J.JCPS.2010.08.004

27. Falk, E. B., Berkman, E. T., & Lieberman, M. D. (2012). From Neural Responses to Population Behavior: Neural Focus Group Predicts Population-Level Media Effects. Psychological Science, 23(5), 439–445. 10.1177/0956797611434964/ASSET/IMAGES/LARGE/10.1177_0956797611434964-FIG2.JPEG

28. Fisher, R. J. (1993). Social Desirability Bias and the Validity of Indirect Questioning. Journal of Consumer Research, 20(2), 303–315. 10.1086/209351

29. Genevsky, A., Tong, L. C., & Knutson, B. (2025). Neuroforecasting reveals generalizable components of choice. PNAS Nexus, 4(2). 10.1093/PNASNEXUS/PGAF029

30. Genevsky, A., Yoon, C., & Knutson, B. (2017). When Brain Beats Behavior: Neuroforecasting Crowdfunding Outcomes. The Journal of Neuroscience, 37(36), 8625–8634. 10.1523/JNEUROSCI.1633-16.2017

31. Guixeres, J., Bigné, E., Azofra, J. M. A., Raya, M. A., Granero, A. C., Hurtado, F. F., & Ornedo, V. N. (2017). Consumer neuroscience-based metrics predict recall, liking and viewing rates in online advertising. Frontiers in Psychology, 8(OCT), 291886. 10.3389/FPSYG.2017.01808/BIBTEX

32. Gupta, R., Kapoor, A. P., & Verma, H. V. (2025). Neuro-insights: A systematic review of neuromarketing perspectives across consumer buying stages. Frontiers in Neuroergonomics, 6, 1542847–1542847. 10.3389/FNRGO.2025.1542847/XML

33. Hakim, A., Golan, I., Yefet, S., & Levy, D. J. (2023). DeePay: Deep learning decodes EEG to predict consumer’s willingness to pay for neuromarketing. Frontiers in Human Neuroscience, 17, 1153413. 10.3389/FNHUM.2023.1153413/BIBTEX

34. Hakim, A., Klorfeld, S., Sela, T., Friedman, D., Shabat-Simon, M., & Levy, D. J. (2018). Pathways to Consumers’ Minds: Using Machine Learning and Multiple EEG Metrics to Increase Preference Prediction Above and Beyond Traditional Measurements. bioRxiv 317073. 10.1101/317073

35. Hakim, A., Klorfeld, S., Sela, T., Friedman, D., Shabat-Simon, M., & Levy, D. J. (2021). Machines learn neuromarketing: Improving preference prediction from self-reports using multiple EEG measures and machine learning. International Journal of Research in Marketing, 38(3), 770–791. 10.1016/J.IJRESMAR.2020.10.005

36. Harris, J. M., Ciorciari, J., & Gountas, J. (2018). Consumer neuroscience for marketing researchers. Journal of Consumer Behaviour, 17(3), 239–252. 10.1002/CB.1710

37. Hasson, U., Nir, Y., Levy, I., Fuhrmann, G., & Malach, R. (2004). Intersubject Synchronization of Cortical Activity during Natural Vision. Science, 303(5664), 1634–1640. 10.1126/SCIENCE.1089506/SUPPL_FILE/HASSON.SOM.PDF

38. Horr, N. K., Han, K., Mousavi, B., & Tang, R. (2022). Neural Signature of Buying Decisions in Real-World Online Shopping Scenarios – An Exploratory Electroencephalography Study Series. Frontiers in Human Neuroscience, 15, 797064. 10.3389/FNHUM.2021.797064/BIBTEX

39. Hsu, M., & Yoon, C. (2015). The neuroscience of consumer choice. *Current Opinion in Behavioral Sciences*, Neuroeconomics, 5, 116–121. 10.1016/j.cobeha.2015.09.005

40. Ingolfsson, T. M., Hersche, M., Wang, X., Kobayashi, N., Cavigelli, L., & Benini, L. (2020). EEG-TCNet: An Accurate Temporal Convolutional Network for Embedded Motor-Imagery Brain-Machine Interfaces. *Conference Proceedings – IEEE International Conference on Systems*, Man and Cybernetics, 2020-October, 2958–2965. 10.1109/SMC42975.2020.9283028

41. Ishtiaque, F., Miya, M. T. I., Mashrur, F. R., Rahman, K. M., Vaidyanathan, R., Anwar, S. F., Sarker, F., Tat, H. H., Hamid, A. B. A., & Mamun, K. A. (2025). Systematic comparison between a research-grade EEG device and a consumer-grade BCI device for predicting consumer preference using an ML framework. Multimedia Tools and Applications, 1–19. 10.1007/S11042-025-20812-3/TABLES/2

42. Johansson, P., Hall, L., Sikström, S., Tärning, B., & Lind, A. (2006). How something can be said about telling more than we can know: On choice blindness and introspection. Consciousness and Cognition, 15(4), 673– 692. 10.1016/J.CONCOG.2006.09.004

43. Karmarkar, U. R., & Plassmann, H. (2017). Consumer Neuroscience: Past, Present, and Future. 10.1177/1094428117730598, 22(1), 174–195. https://doi.org/10.1177/1094428117730598

44. Karmarkar, U. R., & Yoon, C. (2016). Consumer neuroscience: Advances in understanding consumer psychology. *Current Opinion in Psychology*, Consumer Behavior, 10, 160–165. 10.1016/j.copsyc.2016.01.010

45. Khurana, V., Gahalawat, M., Kumar, P., Roy, P. P., Dogra, D. P., Scheme, E., & Soleymani, M. (2021). A Survey on Neuromarketing Using EEG Signals. IEEE Transactions on Cognitive and Developmental Systems, 13(4), 732–749. 10.1109/TCDS.2021.3065200

46. Khushaba, R. N., Greenacre, L., Kodagoda, S., Louviere, J., Burke, S., & Dissanayake, G. (2012). Choice modeling and the brain: A study on the Electroencephalogram (EEG) of preferences. Expert Systems with Applications, 39(16), 12378–12388. 10.1016/J.ESWA.2012.04.084

47. Khushaba, R. N., Wise, C., Kodagoda, S., Louviere, J., Kahn, B. E., & Townsend, C. (2013). Consumer neuroscience: Assessing the brain response to marketing stimuli using electroencephalogram (EEG) and eye tracking. Expert Systems with Applications, 40(9), 3803–3812. 10.1016/J.ESWA.2012.12.095

48. Knutson, B., & Genevsky, A. (2018). Neuroforecasting Aggregate Choice. Current Directions in Psychological Science, 27(2), 110–115. 10.1177/0963721417737877/ASSET/IMAGES/LARGE/10.1177_0963721417737877-FIG1.JPEG

49. Koelstra, S., Mühl, C., Soleymani, M., Lee, J. S., Yazdani, A., Ebrahimi, T., Pun, T., Nijholt, A., & Patras, I. (2012a). DEAP: A database for emotion analysis; Using physiological signals. IEEE Transactions on Affective Computing, 3(1), 18–31. 10.1109/T-AFFC.2011.15

50. Koelstra, S., Mühl, C., Soleymani, M., Lee, J. S., Yazdani, A., Ebrahimi, T., Pun, T., Nijholt, A., & Patras, I. (2012b). DEAP: A database for emotion analysis; Using physiological signals. IEEE Transactions on Affective Computing, 3(1), 18–31. 10.1109/T-AFFC.2011.15

51. Kühn, S., Strelow, E., & Gallinat, J. (2016). Multiple “buy buttons” in the brain: Forecasting chocolate sales at point-of-sale based on functional brain activation using fMRI. NeuroImage, 136, 122–128. 10.1016/J.NEUROIMAGE.2016.05.021

52. Kumar, S., Yadava, M., & Roy, P. P. (2019). Fusion of EEG response and sentiment analysis of products review to predict customer satisfaction. Information Fusion, 52, 41–52. 10.1016/J.INFFUS.2018.11.001

53. Lawhern, V. J., Solon, A. J., Waytowich, N. R., Gordon, S. M., Hung, C. P., & Lance, B. J. (2016). EEGNet: A Compact Convolutional Network for EEG-based Brain-Computer Interfaces. Journal of Neural Engineering, 15(5). 10.1088/1741-2552/aace8c

54. Lin, M. H. (Jenny), Cross, S. N. N., Jones, W. J., & Childers, T. L. (2018). Applying EEG in consumer neuroscience. European Journal of Marketing, 52(1–2), 66–91. 10.1108/EJM-12-2016-0805

55. Luck, S. J. (2014). An Introduction to the Event-Related Potential Technique, second edition. MIT Press. http://www.worldcat.org/title/introduction-to-the-event-related-potential-technique/oclc/861671073

56. McDaniel, Carl., & Gates, Roger. (2016). Marketing research essentials. 12–12.

57. Mcdaniel, S. W., Verille, P., & Madden, C. S. (1985). The Threats to Marketing Research: An Empirical Reappraisal. 10.1177/002224378502200107, 22(1), 74–80. https://doi.org/10.1177/002224378502200107

58. Motoki, K., Suzuki, S., Kawashima, R., & Sugiura, M. (2020). A Combination of Self-Reported Data and Social-Related Neural Measures Forecasts Viral Marketing Success on Social Media. Journal of Interactive Marketing, 52(1), 99–117. 10.1016/j.intmar.2020.06.003

59. Murugappan, M., Murugappan, S., Balaganapathy, B., & Gerard, C. (2014). Wireless EEG signals based Neuromarketing system using Fast Fourier Transform (FFT). Proceedings – 2014 IEEE 10th International Colloquium on Signal Processing and Its Applications, CSPA 2014, 25–30. 10.1109/CSPA.2014.6805714

60. Musallam, Y. K., AlFassam, N. I., Muhammad, G., Amin, S. U., Alsulaiman, M., Abdul, W., Altaheri, H., Bencherif, M. A., & Algabri, M. (2021). Electroencephalography-based motor imagery classification using temporal convolutional network fusion. Biomedical Signal Processing and Control, 69, 102826–102826. 10.1016/J.BSPC.2021.102826

61. Nakagawa, S., & Schielzeth, H. (2013). A general and simple method for obtaining R2 from generalized linear mixed-effects models. Methods in Ecology and Evolution, 4(2), 133–142. 10.1111/j.2041-210x.2012.00261.x

62. Neeley, S. M., & Cronley, M. L. (2004). When Research Participants Don&#146;T Tell It Like It Is: Pinpointing the Effects of Social Desirability Bias Using Self Vs. Indirect-Questioning. ACR North American Advances, NA-31.

63. Newton-Fenner, A., Hewitt, D., Henderson, J., Roberts, H., Mari, T., Gu, Y., Gorelkina, O., Giesbrecht, T., Fallon, N., Roberts, C., & Stancak, A. (2023). Economic value in the Brain: A meta-analysis of willingness-to-pay using the Becker-DeGroot-Marschak auction. PLOS ONE, 18(7), e0286969. 10.1371/journal.pone.0286969

64. Ohme, R., Reykowska, D., Wiener, D., & Choromanska, A. (2009). Analysis of Neurophysiological Reactions to Advertising Stimuli by Means of EEG and Galvanic Skin Response Measures. Journal of Neuroscience, Psychology, and Economics, 2(1), 21–31. 10.1037/A0015462

65. Ohme, R., Reykowska, D., Wiener, D., & Choromanska, A. (2010). Application of frontal EEG asymmetry to advertising research. Journal of Economic Psychology, 31(5), 785–793. 10.1016/J.JOEP.2010.03.008

66. Poulsen, A. T., Kamronn, S., Dmochowski, J., Parra, L. C., & Hansen, L. K. (2017). EEG in the classroom: Synchronised neural recordings during video presentation. Scientific Reports, 7(1), 1–9. 10.1038/SREP43916/FIGURES/5

67. Pratama, B. G., Wibawa, A. D., Wulandari, D. P., & Pratasik, S. (2024). Neuromarketing Study of Purchase Decisions Using Advertising Videos Based on EEG Signal Analysis. Proceedings of the 2024 IEEE International Conference on Industry 4.0, Artificial Intelligence, and Communications Technology, IAICT 2024, 315–320. 10.1109/IAICT62357.2024.10617623

68. Ramsøy, T. Z., Skov, M., Christensen, M. K., & Stahlhut, C. (2018). Frontal brain asymmetry and willingness to pay. Frontiers in Neuroscience, 12(MAR), 283405. 10.3389/FNINS.2018.00138/BIBTEX

69. Ravaja, N., Somervuori, O., & Salminen, M. (2013a). Predicting purchase decision: The role of hemispheric asymmetry over the frontal cortex. Journal of Neuroscience, Psychology, and Economics, 6(1), 1–13. 10.1037/A0029949

70. Ravaja, N., Somervuori, O., & Salminen, M. (2013b). Predicting purchase decision: The role of hemispheric asymmetry over the frontal cortex. Journal of Neuroscience, Psychology, and Economics, 6(1), 1–13. 10.1037/A0029949

71. Schirrmeister, R. T., Springenberg, J. T., Fiederer, L. D. J., Glasstetter, M., Eggensperger, K., Tangermann, M., Hutter, F., Burgard, W., & Ball, T. (2017). Deep learning with convolutional neural networks for EEG decoding and visualization. Human Brain Mapping, 38(11), 5391–5420. 10.1002/HBM.23730

72. Scholz, C., Baek, E. C., O’Donnell, M. B., Kim, H. S., Cappella, J. N., & Falk, E. B. (2017). A neural model of valuation and information virality. Proceedings of the National Academy of Sciences of the United States of America, 114(11), 2881–2886. 10.1073/PNAS.1615259114/SUPPL_FILE/PNAS.201615259SI.PDF

73. Shmueli, G. (2010). To Explain or to Predict? Statistical Science, 25(3), 289–310. 10.1214/10-STS330

74. Smidts, A., Hsu, M., Sanfey, A. G., Boksem, M. A. S., Ebstein, R. B., Huettel, S. A., Kable, J. W., Karmarkar, U. R., Kitayama, S., Knutson, B., Liberzon, I., Lohrenz, T., Stallen, M., & Yoon, C. (2014). Advancing consumer neuroscience. Marketing Letters, 25(3), 257–267. 10.1007/s11002-014-9306-1

75. Smith, M. E., & Gevins, A. (2004). Attention and Brain Activity While Watching Television: Components of Viewer Engagement. Media Psychology, 6(3), 285–305. 10.1207/S1532785XMEP0603_3

76. Sutton, S. K., & Davidson, R. J. (2000). Prefrontal brain electrical asymmetry predicts the evaluation of affective stimuli. Neuropsychologia, 38(13), 1723–1733. 10.1016/S0028-3932(00)00076-2

77. Telpaz, A., Webb, R., & Levy, D. J. (2015). Using EEG to predict consumers’ future choices. Journal of Marketing Research, 52(4), 511–529. 10.1509/JMR.13.0564/ASSET/IMAGES/LARGE/10.1509_JMR.13.0564-FIG8.JPEG

78. Varga, M., Tusche, A., Albuquerque, P., Gier, N., Weber, B., & Plassmann, H. (2021). Marketing Science Institute Working Paper Series 2021 Predicting Sales of New Consumer Packaged Products with fMRI, Behavioral, Survey and Market Data “Predicting Sales of New Consumer Packaged Products with fMRI, Behavioral, Survey and Market Data” © 2021 PREDICTING SALES OF NEW CONSUMER PACKAGED PRODUCTS WITH fMRI, BEHAVIORAL, SURVEY, AND MARKET DATA.

79. Vecchiato, G., Astolfi, L., Fallani, F. D. V., Cincotti, F., Mattia, D., Salinari, S., Soranzo, R., & Babiloni, F. (2010). Changes in brain activity during the observation of TV commercials by using EEG, GSR and HR measurements. Brain Topography, 23(2), 165–179. 10.1007/S10548-009-0127-0/FIGURES/6

80. Vecchiato, G., Cherubino, P., Maglione, A. G., Ezquierro, M. T. H., Marinozzi, F., Bini, F., Trettel, A., & Babiloni, F. (2014). How to Measure Cerebral Correlates of Emotions in Marketing Relevant Tasks. Cognitive Computation, 6(4), 856–871. 10.1007/s12559-014-9304-x

81. Vecchiato, G., Toppi, J., Astolfi, L., De Vico Fallani, F., Cincotti, F., Mattia, D., Bez, F., & Babiloni, F. (2011). Spectral EEG frontal asymmetries correlate with the experienced pleasantness of TV commercial advertisements. Medical and Biological Engineering and Computing, 49(5), 579–583. 10.1007/S11517-011-0747-X/FIGURES/2

82. Venkatraman, V., Dimoka, A., Pavlou, P. A., Vo, K., Hampton, W., Bollinger, B., Hershfield, H. E., Ishihara, M., & Winer, R. S. (2015). Predicting Advertising success beyond Traditional Measures: New Insights from Neurophysiological Methods and Market Response Modeling: Journal of Marketing Research, 52(4), 436–452. 10.1509/JMR.13.0593

83. Wang, R. W. Y., Chang, Y. C., & Chuang, S. W. (2016). EEG Spectral Dynamics of Video Commercials: Impact of the Narrative on the Branding Product Preference. Scientific Reports 2016 6:1, 6(1), 1–11. 10.1038/srep36487

84. Wilkinson, D., & Birmingham, P. (2003). Using Research Instruments: A Guide for Researchers. Psychology Press.

85. Xu, Z., & Liu, S. (2024). Decoding consumer purchase decisions: Exploring the predictive power of EEG features in online shopping environments using machine learning. Humanities and Social Sciences Communications, 11(1), 1–13. 10.1057/S41599-024-03691-1;SUBJMETA=4000,4001,4014,4045;KWRD=BUSINESS+AND+MANAGEMENT,SCIENCE

86. Yadava, M., Kumar, P., Saini, R., Roy, P. P., & Prosad Dogra, D. (2017). Analysis of EEG signals and its application to neuromarketing. Multimedia Tools and Applications, 76(18), 19087–19111. 10.1007/S11042-017-4580-6/TABLES/4

87. Yarkoni, T., & Westfall, J. (2017). Choosing Prediction Over Explanation in Psychology: Lessons From Machine Learning. Perspectives on Psychological Science, 12(6), 1100–1122. 10.1177/1745691617693393

88. Yu, W., Sun, Z., Xu, T., & Ma, Q. (2018). Things Become Appealing When I Win: Neural Evidence of the Influence of Competition Outcomes on Brand Preference. Frontiers in Neuroscience, 12, 411509. 10.3389/FNINS.2018.00779/BIBTEX

89. Zeng, L., Lin, M., Xiao, K., Wang, J., & Zhou, H. (2022). Like/Dislike Prediction for Sport Shoes With Electroencephalography: An Application of Neuromarketing. Frontiers in Human Neuroscience, 15, 793952. 10.3389/FNHUM.2021.793952/BIBTEX

90. Zhao, T., Gois, F. N. B., Marques, J. A. L., dos Santos Silva, B. R., Cortez, P., Jagatheesaperumal, S. K., & de Albuquerque, V. H. C. (2025). AutoML-based EEG signal analysis in neuro-marketing classification using biclustering method. Applied Soft Computing, 181, 113412–113412. 10.1016/J.ASOC.2025.113412

