## Supplementary material for "Predicting the immediate and subsequent effects of commercials on product valuation using EEG and deep learning": full supplemetary file

**Web Appendices**

These materials have been supplied by the authors to aid in the understanding of their paper. The AMA is sharing these materials at the request of the authors.

### **Web Appendix A: Stimuli**

| **Category** | **Product** |
| --- | --- |
| Beverages | Cafeelite; Chocolatemilk; CocaCola2L; FuzeTeaGreen; Neviot; Wissotzky |
| Breakfast | Branflakes; Branflakesextra; ChocolateSpread; Quaker |
| Dry Goods | Filtuna; Heinz; Kikkoman; Prince; YadMordechai |
| Hygiene | Colgate; GelDeLin; Sano99; Zipperbigbag; Zippersmallbag |
| Snacks | Fitness; Oreo; Parachocolate; Splendid; Tapuchips Baracke |

**Table W1**. list of products used in the experiment, divided by category

### **Web Appendix B: EEG Features – Spectral Power**

|  | Coefficient | SE | z | p | Lower CI | Upper CI |
| --- | --- | --- | --- | --- | --- | --- |
| Intercept | **0.4177** | **0.013** | **31.423** | **0** | **0.392** | **0.444** |
| Fp2_gamma | **0.0194** | **0.008** | **2.499** | **0.012** | **0.004** | **0.035** |
| Fz_gamma | **-0.0079** | **0.002** | **-3.305** | **0.001** | **-0.013** | **-0.003** |
| Pz_gamma | **0.0069** | **0.003** | **2.35** | **0.019** | **0.001** | **0.013** |
| F7_gamma | -0.0083 | 0.006 | -1.376 | 0.169 | -0.02 | 0.004 |
| Fpz_theta | **-0.0014** | **0.001** | **-2.555** | **0.011** | **-0.002** | **0** |
| Fp1_beta | -0.0159 | 0.009 | -1.724 | 0.085 | -0.034 | 0.002 |
| Fpz_gamma | -0.0149 | 0.009 | -1.716 | 0.086 | -0.032 | 0.002 |
| Fp1_delta | 0.0002 | 0 | 1.6 | 0.11 | -4.2E-05 | 0 |
| F7_delta | **-0.0006** | **0** | **-2.726** | **0.006** | **-0.001** | **0** |
| F7_beta | **0.0142** | **0.007** | **2.087** | **0.037** | **0.001** | **0.028** |
| Fz_theta | -0.0031 | 0.002 | -1.66 | 0.097 | -0.007 | 0.001 |
| Pz_theta | 0.0078 | 0.005 | 1.696 | 0.09 | -0.001 | 0.017 |
| Fp2_theta | 0.0011 | 0.001 | 1.387 | 0.165 | 0 | 0.003 |
| Cz_beta | -0.009 | 0.006 | -1.61 | 0.107 | -0.02 | 0.002 |
| Fp2_alpha | 0.0053 | 0.006 | 0.967 | 0.333 | -0.005 | 0.016 |
| F8_beta | -0.0022 | 0.007 | -0.293 | 0.77 | -0.017 | 0.012 |
| Fp2_delta | -0.0001 | 0 | -1.042 | 0.298 | 0 | 0 |
| Cz_delta | 0.001 | 0.001 | 1.902 | 0.057 | -3.1E-05 | 0.002 |
| F7_alpha | 0.0052 | 0.005 | 1.012 | 0.312 | -0.005 | 0.015 |
| Fz_alpha | -0.0039 | 0.003 | -1.2 | 0.23 | -0.01 | 0.002 |
| F7_theta | 0.0022 | 0.002 | 1.161 | 0.245 | -0.001 | 0.006 |
| Fp1_alpha | -0.0039 | 0.006 | -0.689 | 0.491 | -0.015 | 0.007 |
| Cz_theta | -0.0045 | 0.005 | -0.958 | 0.338 | -0.014 | 0.005 |
| F8_gamma | 0.0047 | 0.007 | 0.722 | 0.47 | -0.008 | 0.017 |
| F8_delta | -0.0002 | 0 | -0.509 | 0.611 | -0.001 | 0.001 |
| F8_alpha | 0.0041 | 0.009 | 0.473 | 0.636 | -0.013 | 0.021 |
| Fp2_beta | 0.0092 | 0.009 | 0.997 | 0.319 | -0.009 | 0.027 |
| F8_theta | 0.0012 | 0.004 | 0.347 | 0.729 | -0.006 | 0.008 |
| Cz_alpha | 0.0041 | 0.004 | 1.129 | 0.259 | -0.003 | 0.011 |
| Pz_alpha | -0.0025 | 0.002 | -1.033 | 0.301 | -0.007 | 0.002 |

**Table W2.** Relaxed elastic net model results of the relationship between spectral power and WTP in Part 1. Each row represents a predictor comprised of a frequency band and an electrode. In bold are the significant coefficients.

|  | Coefficient | SE | z | p | Lower CI | Upper CI |
| --- | --- | --- | --- | --- | --- | --- |
| Intercept | **0.4612** | **0.016** | **28.174** | **0** | **0.429** | **0.493** |
| Fp1_gamma | **0.0212** | **0.009** | **2.419** | **0.016** | **0.004** | **0.038** |
| Fpz_beta | 0.0141 | 0.017 | 0.832 | 0.405 | -0.019 | 0.047 |
| Fpz_gamma | -0.016 | 0.011 | -1.471 | 0.141 | -0.037 | 0.005 |
| Fp1_alpha | -0.016 | 0.011 | -1.461 | 0.144 | -0.037 | 0.005 |
| F8_gamma | -0.01 | 0.009 | -1.089 | 0.276 | -0.028 | 0.008 |
| Fpz_alpha | 0.0189 | 0.015 | 1.303 | 0.193 | -0.01 | 0.047 |
| Fz_beta | -0.0048 | 0.005 | -1.033 | 0.302 | -0.014 | 0.004 |
| Cz_theta | **-0.0148** | **0.005** | **-3.191** | **0.001** | **-0.024** | **-0.006** |
| Pz_gamma | 0.0105 | 0.006 | 1.77 | 0.077 | -0.001 | 0.022 |
| Pz_theta | **0.0136** | **0.004** | **3.637** | **0** | **0.006** | **0.021** |
| Fp1_delta | **0.0005** | **0** | **3.043** | **0.002** | **0** | **0.001** |
| Pz_beta | 0.0061 | 0.005 | 1.292 | 0.196 | -0.003 | 0.015 |
| F7_beta | **-0.0078** | **0.003** | **-2.647** | **0.008** | **-0.014** | **-0.002** |
| Fz_gamma | **-0.0082** | **0.004** | **-2.336** | **0.019** | **-0.015** | **-0.001** |
| Fp2_theta | 0.0038 | 0.002 | 1.94 | 0.052 | 0 | 0.008 |
| Fz_delta | -0.0009 | 0.001 | -1.603 | 0.109 | -0.002 | 0 |
| Fp2_beta | -0.0075 | 0.017 | -0.452 | 0.651 | -0.04 | 0.025 |
| Fp2_alpha | -0.0139 | 0.01 | -1.45 | 0.147 | -0.033 | 0.005 |
| Fpz_delta | -0.0002 | 0 | -0.551 | 0.581 | -0.001 | 0 |
| Fp2_gamma | -0.0036 | 0.009 | -0.391 | 0.696 | -0.022 | 0.015 |
| F7_gamma | 0.0055 | 0.008 | 0.69 | 0.49 | -0.01 | 0.021 |
| F7_alpha | **0.0094** | **0.004** | **2.116** | **0.034** | **0.001** | **0.018** |
| Cz_delta | 0.0011 | 0.001 | 1.254 | 0.21 | -0.001 | 0.003 |
| Fpz_theta | -0.0008 | 0.002 | -0.36 | 0.719 | -0.005 | 0.003 |
| F8_theta | 0.0019 | 0.004 | 0.477 | 0.633 | -0.006 | 0.01 |
| Pz_alpha | 0.0012 | 0.003 | 0.475 | 0.635 | -0.004 | 0.006 |
| Fz_alpha | -0.0016 | 0.004 | -0.384 | 0.701 | -0.01 | 0.007 |
| F7_theta | 0.0014 | 0.002 | 0.686 | 0.493 | -0.003 | 0.005 |
| F8_alpha | 0.0037 | 0.007 | 0.499 | 0.618 | -0.011 | 0.018 |
| Fz_theta2 | -0.0009 | 0.003 | -0.352 | 0.725 | -0.006 | 0.004 |
| Pz_delta | -0.0005 | 0.001 | -0.941 | 0.347 | -0.002 | 0.001 |
| Cz_beta | 0.0005 | 0.004 | 0.144 | 0.885 | -0.007 | 0.008 |
| F8_delta | 9.66E-05 | 0 | 0.224 | 0.822 | -0.001 | 0.001 |
| Cz_alpha | -0.0016 | 0.004 | -0.391 | 0.696 | -0.01 | 0.007 |
| Fp2_delta2 | -0.0003 | 0 | -0.985 | 0.325 | -0.001 | 0 |
| Fp1_theta | -0.002 | 0.002 | -1.184 | 0.237 | -0.005 | 0.001 |
| F7_delta | -0.0001 | 0 | -0.518 | 0.604 | -0.001 | 0 |

**Table W3**. Relaxed elastic net model results of the relationship between spectral power and WTP in Part 2. Each row represents a predictor comprised of a frequency band and an electrode. In bold are the significant coefficients.

|  | Coefficient | SE | z | p | Lower CI | Upper CI |
| --- | --- | --- | --- | --- | --- | --- |
| Intercept | **0.5108** | **0.014** | **36.545** | **0** | **0.483** | **0.538** |
| Cz_gamma | **-0.0165** | **0.008** | **-2.193** | **0.028** | **-0.031** | **-0.002** |
| Fpz_gamma | 0.0085 | 0.009 | 0.987 | 0.324 | -0.008 | 0.025 |
| Fp1_gamma | 0.0066 | 0.008 | 0.872 | 0.383 | -0.008 | 0.021 |
| Fz_delta | **-0.0013** | **0** | **-4.256** | **0** | **-0.002** | **-0.001** |
| F8_alpha | **-0.014** | **0.007** | **-2.072** | **0.038** | **-0.027** | **-0.001** |
| Fp1_alpha | -0.0104 | 0.007 | -1.581 | 0.114 | -0.023 | 0.002 |
| Cz_delta | **0.0017** | **0.001** | **2.91** | **0.004** | **0.001** | **0.003** |
| Fz_gamma | -0.003 | 0.003 | -0.992 | 0.321 | -0.009 | 0.003 |
| F7_gamma | -0.0056 | 0.007 | -0.831 | 0.406 | -0.019 | 0.008 |
| Cz_theta | -0.0053 | 0.003 | -1.845 | 0.065 | -0.011 | 0 |
| Fp2_alpha | 0.0004 | 0.011 | 0.038 | 0.969 | -0.02 | 0.021 |
| Fpz_alpha | 0.0082 | 0.013 | 0.621 | 0.535 | -0.018 | 0.034 |
| F8_gamma | 0.0056 | 0.009 | 0.659 | 0.51 | -0.011 | 0.022 |
| Fpz_delta | 0.0001 | 0 | 0.386 | 0.699 | 0 | 0.001 |
| F7_alpha | 0.0086 | 0.005 | 1.897 | 0.058 | 0 | 0.018 |
| Pz_gamma | 0.0041 | 0.005 | 0.893 | 0.372 | -0.005 | 0.013 |
| F8_delta | -0.0005 | 0 | -1.358 | 0.175 | -0.001 | 0 |
| F7_beta | -0.0024 | 0.002 | -0.979 | 0.327 | -0.007 | 0.002 |
| Pz_alpha | -0.0024 | 0.002 | -1.06 | 0.289 | -0.007 | 0.002 |
| Pz_theta | 0.0037 | 0.003 | 1.159 | 0.247 | -0.003 | 0.01 |
| F8_theta | 0.0023 | 0.003 | 0.723 | 0.47 | -0.004 | 0.009 |
| F7_theta | -0.0017 | 0.002 | -0.98 | 0.327 | -0.005 | 0.002 |
| F7_delta | 0.0002 | 0 | 0.869 | 0.385 | 0 | 0.001 |
| Fp2_delta | 0.00005 | 0 | 0.173 | 0.862 | -0.001 | 0.001 |
| Fpz_theta | 0.0006 | 0.002 | 0.287 | 0.774 | -0.003 | 0.004 |
| Fp2_theta | 0.0004 | 0.002 | 0.181 | 0.856 | -0.004 | 0.004 |
| Cz_alpha | 0.0008 | 0.003 | 0.232 | 0.816 | -0.006 | 0.007 |
| Pz_delta | 0.0003 | 0 | 0.779 | 0.436 | 0 | 0.001 |
| Fp2_beta | 0.0014 | 0.003 | 0.489 | 0.625 | -0.004 | 0.007 |

**Table W4**. Relaxed elastic net model results of the relationship between spectral power and liking in Part 2. Each row represents a predictor comprised of a frequency band and an electrode. In bold are the significant coefficients.

|  | Coefficient | SE | z | p | Lower CI | Upper CI |
| --- | --- | --- | --- | --- | --- | --- |
| Intercept | **0.437** | **0.017** | **25.997** | **0** | **0.404** | **0.47** |
| Fz_theta | -0.0033 | 0.002 | -1.426 | 0.154 | -0.008 | 0.001 |
| F7_alpha | **0.0171** | **0.007** | **2.363** | **0.018** | **0.003** | **0.031** |
| F7_delta | **-0.0011** | **0** | **-2.464** | **0.014** | **-0.002** | **0** |
| Pz_theta | **0.0075** | **0.004** | **1.973** | **0.048** | **5.13E-05** | **0.015** |
| Fz_alpha | -0.0084 | 0.006 | -1.355 | 0.175 | -0.02 | 0.004 |
| Cz_alpha | -0.0017 | 0.006 | -0.297 | 0.766 | -0.013 | 0.009 |
| Cz_delta | 0.0009 | 0.001 | 0.739 | 0.46 | -0.001 | 0.003 |
| Pz_beta | 0.001 | 0.002 | 0.413 | 0.679 | -0.004 | 0.006 |
| Fp1_delta | 0.0002 | 0 | 1.409 | 0.159 | -0.00009 | 0.001 |
| Fpz_theta | -0.0007 | 0.001 | -0.891 | 0.373 | -0.002 | 0.001 |
| Pz_alpha | -0.0039 | 0.004 | -1.033 | 0.302 | -0.011 | 0.004 |

**Table W5**. Relaxed elastic net model results of the relationship between spectral power in Part 1 and WTP in Part 3. Each row represents a predictor comprised of a frequency band and an electrode. In bold are the significant coefficients.

### **Web Appendix C: EEG Features – Inter Subject Correlation**

|  | Coefficient | SE | z | p | Lower CI | Upper CI |
| --- | --- | --- | --- | --- | --- | --- |
| Intercept | **37.184** | **2.1** | **17.703** | **0.000** | **33.067** | **41.3** |
| theta | 2.145 | 3.378 | 0.635 | 0.956 | -4.475 | 8.765 |
| delta | -1.144 | 3.169 | -0.361 | 0.956 | -7.355 | 5.067 |
| alpha | -0.176 | 3.235 | -0.055 | 0.956 | -6.517 | 6.165 |
| beta | -4.18 | 3.306 | -1.264 | 0.956 | -10.661 | 2.301 |
| gamma | 0.455 | 1.727 | 0.258 | 0.956 | -2.939 | 3.829 |

**Table W6**. Mixed effects model results of the relationship between ISC and WTP in Part 1. Each row represents a frequency band. In bold are the significant coefficients. Predictor coefficients after an FDR correction

|  |  | Coefficient | SE | z | p | Lower CI | Upper CI |
| --- | --- | --- | --- | --- | --- | --- | --- |
| Intercept | **WTP** | **38.845** | **1.534** | **25.316** | **0.000** | **35.838** | **41.852** |
|  | **Liking** | **3.215** | **0.197** | **16.303** | **0.000** | **2.828** | **3.601** |
| theta | WTP | -2.234 | 2.737 | -0.816 | 0.517 | -7.598 | 3.13 |
|  | Liking | 0.303 | 0.375 | 0.808 | 0.989 | -0.432 | 1.037 |
| delta | WTP | 3.528 | 2.581 | 1.367 | 0.429 | -1.53 | 8.585 |
|  | Liking | -0.107 | 0.353 | -0.304 | 0.989 | -0.799 | 0.585 |
| alpha | WTP | 0.251 | 1.972 | 0.128 | 0.898 | -3.613 | 4.116 |
|  | Liking | -0.004 | 0.268 | -0.013 | 0.989 | -0.528 | 0.521 |
| beta | WTP | -2.318 | 2.619 | -0.885 | 0.518 | -7.45 | 2.815 |
|  | Liking | 0.030 | 0.358 | 0.084 | 0.989 | -0.672 | 0.733 |
| gamma | WTP | 2.961 | 1.845 | 1.605 | 0.429 | -0.656 | 6.577 |
|  | **Liking** | **0.679** | **0.248** | **2.728** | **0.031** | **0.19** | **1.16** |

**Table W7**. Mixed-effects models result of the relationship between ISC and WTP/Liking in Part 2. Each row represents a frequency band. In bold are the significant coefficients. Predictors are after an FDR correction.

|  | Coefficient | SE | z | p | Lower CI | Upper CI |
| --- | --- | --- | --- | --- | --- | --- |
| Intercept | **52.047** | **3.848** | **13.526** | **0.000** | **44.505** | **59.588** |
| theta | 9.92 | 8.224 | 1.207 | 0.455 | -6.196 | 26.04 |
| delta | -22.804 | 9.145 | -2.494 | 0.063 | -40.728 | -4.88 |
| alpha | -11.684 | 10.671 | -1.095 | 0.455 | -32.599 | 9.231 |
| beta | -6.344 | 9.308 | -0.682 | 0.495 | -24.587 | 11.90 |
| gamma | -3.520 | 4.115 | -0.855 | 0.490 | -11.585 | 4.546 |

**Table W8.** Mixed-effects models result of the relationship between ISC in Part 1 and WTP in Part 3. Each row represents a frequency band. In bold are the significant coefficients. Predictors are after an FDR correction.

### **Web Appendix D: EEG Features – Frontal Asymmetry**

|  | Coefficient | SE | z | p | Lower CI | Upper CI |
| --- | --- | --- | --- | --- | --- | --- |
| Intercept | **35.455** | **0.553** | **64.163** | **0.000** | **34.372** | **36.538** |
| theta | -0.607 | 0.706 | -0.861 | 0.389 | -1.991 | 0.776 |
| delta | 0.299 | 0.462 | 0.648 | 0.517 | -0.606 | 1.204 |
| alpha | **-1.433** | **0.588** | **-2.569** | **0.017** | **-2.526** | **-0.34** |
| beta | **2.286** | **0.733** | **3.119** | **0.004** | **0.849** | **3.724** |
| gamma | **-2.299** | **0.506** | **-4.544** | **0.000** | **-3.291** | **-1.308** |

**Table W9**. Mixed effects model results of the relationship between FA and WTP in Part 1. Each row represents a frequency band. In bold are the significant coefficients. Predictor coefficients after an FDR correction.

|  |  | Coefficient | SE | z | p | Lower CI | Upper CI |
| --- | --- | --- | --- | --- | --- | --- | --- |
| Intercept | **WTP** | **31.181** | **0.641** | **61.094** | **0.000** | **37.924** | **40.438** |
|  | **Liking** | **3.303** | **0.052** | **63.267** | **0.000** | **3.201** | **3.406** |
| theta | WTP | 0.287 | 0.641 | 0.448 | 0.748 | -0.969 | 1.543 |
|  | Liking | -0.17 | 0.086 | -1.972 | 0.242 | -0.34 | -0.001 |
| delta | WTP | -0.452 | 0.403 | -1.123 | 0.748 | -1.242 | 0.337 |
|  | Liking | -0.016 | 0.055 | -0.289 | 0.772 | -0.123 | 0.091 |
| alpha | WTP | -0.364 | 0.521 | -0.7 | 0.748 | -1.385 | 0.656 |
|  | Liking | -0.072 | 0.07 | -1.042 | 0.608 | -0.209 | 0.064 |
| beta | WTP | 0.229 | 0.714 | 0.32 | 0.748 | -1.17 | -1.628 |
|  | Liking | -0.08 | 0.091 | -0.876 | 0.608 | -0.257 | 0.098 |
| gamma | WTP | -0.218 | 0.58 | -0.375 | 0.748 | -1.355 | 0.919 |
|  | Liking | 0.047 | 0.068 | 0.696 | 0.608 | -0.085 | 0.18 |

**Table W10**. Regression results of the relationship between FA and WTP/Liking. Each row represents a frequency band. In bold are the significant coefficients after an FDR correction.

|  | Coefficient | SE | z | p | Lower CI | Upper CI |
| --- | --- | --- | --- | --- | --- | --- |
| Intercept | **34.539** | **0.894** | **38.632** | **0.000** | **32.787** | **36.291** |
| theta | -1.403 | 1.439 | -0.975 | 0.549 | -4.224 | 1.417 |
| delta | 0.114 | 0.825 | 0.138 | 0.989 | -1.503 | 1.731 |
| alpha | -3.622 | 1.760 | -2.058 | 0.098 | -7.070 | -0.173 |
| beta | -0.022 | 1.711 | -0.013 | 0.989 | -3.376 | 3.332 |
| gamma | **-2.596** | **0.977** | **-2.657** | **0.039** | **-4.511** | **-0.681** |

**Table W11**. Mixed effects model results of the relationship between FA in Part 1 and WTP in Part 3. Each row represents a frequency band. In bold are the significant coefficients. Predictor coefficients after an FDR correction.

### **Web Appendix E: EEG Features – All Features Combined**

|  | Coefficient | SE | z | p | Lower CI | Upper CI |
| --- | --- | --- | --- | --- | --- | --- |
| Intercept | **34.8914** | **1.654** | **21.102** | **0.000** | **31.651** | **38.132** |
| Fpz_theta | -0.0439 | 0.034 | -1.307 | 0.191 | -0.110 | 0.022 |
| beta_asymmetry | -0.8053 | 2.100 | -0.383 | 0.701 | -4.922 | 3.311 |
| delta_Asymmetry | 0.7509 | 1.070 | 0.702 | 0.483 | -1.346 | 2.848 |
| theta_Asymmetry | **-3.7195** | **1.652** | **-2.251** | **0.024** | **-6.958** | **-0.481** |

***Table W12.*** *Relaxed elastic net model results of the relationship between spectral power, ISC, FA and WTP in Part 1. Each row represents a predictor comprised of either a frequency band and an electrode from the spectral power analysis, or a frequency band and ISC or Asymmetry, from the ISC and FA analyses, respectively. In bold are the significant coefficients.*

|  | Coefficient | SE | z | p | Lower CI | Upper CI |
| --- | --- | --- | --- | --- | --- | --- |
| Intercept | **0.3769** | **0.045** | **8.454** | **0.000** | **0.290** | **0.464** |
| Fz_beta | **-0.0125** | **0.003** | **-3.750** | **0.000** | **-0.019** | **-0.006** |
| Fp2_beta | 0.0057 | 0.008 | 0.700 | 0.484 | -0.010 | 0.022 |
| gamma_Asymmetry | -0.0556 | 0.030 | -1.864 | 0.062 | -0.114 | 0.003 |
| delta_Asymmetry | **-0.0303** | **0.012** | **-2.491** | **0.013** | **-0.054** | **-0.006** |
| Fz_delta | **-0.0014** | **0.001** | **-2.240** | **0.025** | **-0.003** | **-0.000** |
| Pz_theta | -0.0065 | 0.004 | -1.642 | 0.101 | -0.014 | 0.001 |
| Fpz_delta | **0.0002** | **9.8e-05** | **2.329** | **0.020** | **3.62e-05** | **0.000** |
| beta_Asymmetry | 0.0613 | 0.036 | 1.693 | 0.091 | -0.010 | 0.132 |
| F8_alpha | 0.0134 | 0.007 | 1.902 | 0.057 | -0.000 | 0.027 |
| F8_delta | -0.0005 | 0.000 | -1.047 | 0.295 | -0.001 | 0.000 |
| Fz_theta | -0.0016 | 0.002 | -0.643 | 0.520 | -0.006 | 0.003 |
| Cz_alpha | 0.0066 | 0.005 | 1.406 | 0.160 | -0.003 | 0.016 |
| F7_beta | 0.0024 | 0.002 | 0.988 | 0.323 | -0.002 | 0.007 |
| delta_isc | **0.1750** | **0.075** | **2.338** | **0.019** | **0.028** | **0.322** |
| Fp1_beta | 0.0027 | 0.007 | 0.393 | 0.694 | -0.011 | 0.016 |
| Fz_gamma | -0.0009 | 0.002 | -0.482 | 0.630 | -0.004 | 0.003 |
| alpha_isc | 0.1153 | 0.074 | 1.563 | 0.118 | -0.029 | 0.260 |
| Cz_delta | 0.0013 | 0.001 | 1.704 | 0.088 | -0.000 | 0.003 |
| F7_alpha | 0.0007 | 0.005 | 0.145 | 0.885 | -0.008 | 0.010 |
| Pz_alpha | -0.0039 | 0.003 | -1.233 | 0.218 | -0.010 | 0.002 |
| alpha_Asymmetry | -0.0038 | 0.021 | -0.178 | 0.859 | -0.046 | 0.038 |
| theta_isc | -0.0961 | 0.083 | -1.156 | 0.248 | -0.259 | 0.067 |

***Table W13.*** *Relaxed elastic net model results of the relationship between spectral power, ISC, FA and WTP in Part 2. Each row represents a predictor comprised of either a frequency band and an electrode from the spectral power analysis, or a frequency band and ISC or Asymmetry, from the ISC and FA analyses, respectively. In bold are the significant coefficients.*

|  | Coefficient | SE | z | p | Lower CI | Upper CI |
| --- | --- | --- | --- | --- | --- | --- |
| Intercept | **0.3796** | **0.039** | **9.667** | **0.000** | **0.303** | **0.457** |
| Fz_beta | **-0.0085** | **0.004** | **-2.201** | **0.028** | **-0.016** | **-0.001** |
| Fp2_beta | 0.0046 | 0.004 | 1.249 | 0.212 | -0.003 | 0.012 |
| gamma_Asymmetry | 0.0319 | 0.028 | 1.124 | 0.261 | -0.024 | 0.087 |
| theta_Asymmetry | -0.0241 | 0.019 | -1.291 | 0.197 | -0.061 | 0.012 |
| beta_Asymmetry | -0.0413 | 0.028 | -1.478 | 0.140 | -0.096 | 0.013 |
| Cz_alpha | -0.0044 | 0.006 | -0.774 | 0.439 | -0.016 | 0.007 |
| F7_theta | **0.0040** | **0.002** | **2.202** | **0.028** | **0.000** | **0.007** |
| Cz_delta | **0.0012** | **0.001** | **2.421** | **0.015** | **0.000** | **0.002** |
| alpha_Asymmetry | -0.0256 | 0.021 | -1.210 | 0.226 | -0.067 | 0.016 |
| F7_beta | 0.0032 | 0.002 | 1.386 | 0.166 | -0.001 | 0.008 |
| Fz_gamma | -0.0039 | 0.003 | -1.400 | 0.161 | -0.009 | 0.002 |
| Fp1_alpha | -0.0020 | 0.005 | -0.404 | 0.686 | -0.012 | 0.008 |
| theta_isc | 0.1008 | 0.064 | 1.573 | 0.116 | -0.025 | 0.226 |
| gamma_isc | **0.1599** | **0.058** | **2.753** | **0.006** | **0.046** | **0.274** |
| Fpz_theta | 0.0009 | 0.001 | 1.305 | 0.192 | -0.000 | 0.002 |
| Pz_alpha | -0.0008 | 0.002 | -0.333 | 0.739 | -0.006 | 0.004 |
| F8_delta | -0.0003 | 0.001 | -0.518 | 0.605 | -0.002 | 0.001 |
| F8_theta | 0.0022 | 0.004 | 0.581 | 0.561 | -0.005 | 0.010 |
| F7_delta | **-0.0004** | **0.000** | **-2.093** | **0.036** | **-0.001** | **-2.83e-05** |
| Fz_alpha | -0.0016 | 0.004 | -0.360 | 0.719 | -0.010 | 0.007 |
| Fp1_gamma | 0.0034 | 0.003 | 1.302 | 0.193 | -0.002 | 0.008 |
| alpha_isc | 0.0075 | 0.067 | 0.113 | 0.910 | -0.123 | 0.138 |
| Fp2_alpha | -0.0032 | 0.006 | -0.525 | 0.600 | -0.015 | 0.009 |

***Table W14.*** *Relaxed elastic net model results of the relationship between spectral power, ISC, FA and Liking in Part 2. Each row represents a predictor comprised of either a frequency band and an electrode from the spectral power analysis, or a frequency band and ISC or Asymmetry, from the ISC and FA analyses, respectively. In bold are the significant coefficients.*

|  | Coefficient | SE | z | p | Lower CI | Upper CI |
| --- | --- | --- | --- | --- | --- | --- |
| Intercept | **42.8628** | **4.289** | **9.993** | **0.000** | **34.456** | **51.270** |
| F7_delta | -0.0499 | 0.046 | -1.090 | 0.276 | -0.140 | 0.040 |
| Fpz_delta | 0.0048 | 0.023 | 0.210 | 0.833 | -0.040 | 0.049 |
| Fz_delta | -0.0520 | 0.078 | -0.669 | 0.504 | -0.204 | 0.100 |
| Cz_delta | 0.0337 | 0.211 | 0.159 | 0.873 | -0.380 | 0.448 |
| Fpz_theta | -0.0596 | 0.097 | -0.616 | 0.538 | -0.249 | 0.130 |
| Fz_theta | -0.2202 | 0.305 | -0.723 | 0.470 | -0.818 | 0.377 |
| Delta_isc | -12.8760 | 7.695 | -1.673 | 0.094 | -27.958 | 2.206 |
| theta_Asymmetry | -2.0808 | 2.639 | -0.788 | 0.430 | -7.254 | 3.092 |
| alpha_ Asymmetry | -4.3044 | 2.946 | -1.461 | 0.144 | -10.079 | 1.470 |
| beta_ Asymmetry | 0.1713 | 3.582 | 0.048 | 0.962 | -6.849 | 7.191 |
| gamma_Asymmetry | -2.5593 | 2.337 | -1.095 | 0.273 | -7.139 | 2.020 |

***Table W15.*** *Relaxed elastic net model results of the relationship between spectral power, ISC and FA in Part 1 and WTP in Part 3. Each row represents a predictor comprised of either a frequency band and an electrode from the spectral power analysis, or a frequency band and ISC or Asymmetry, from the ISC and FA analyses, respectively. In bold are the significant coefficients.*

### **Web Appendix F: Neural Network**

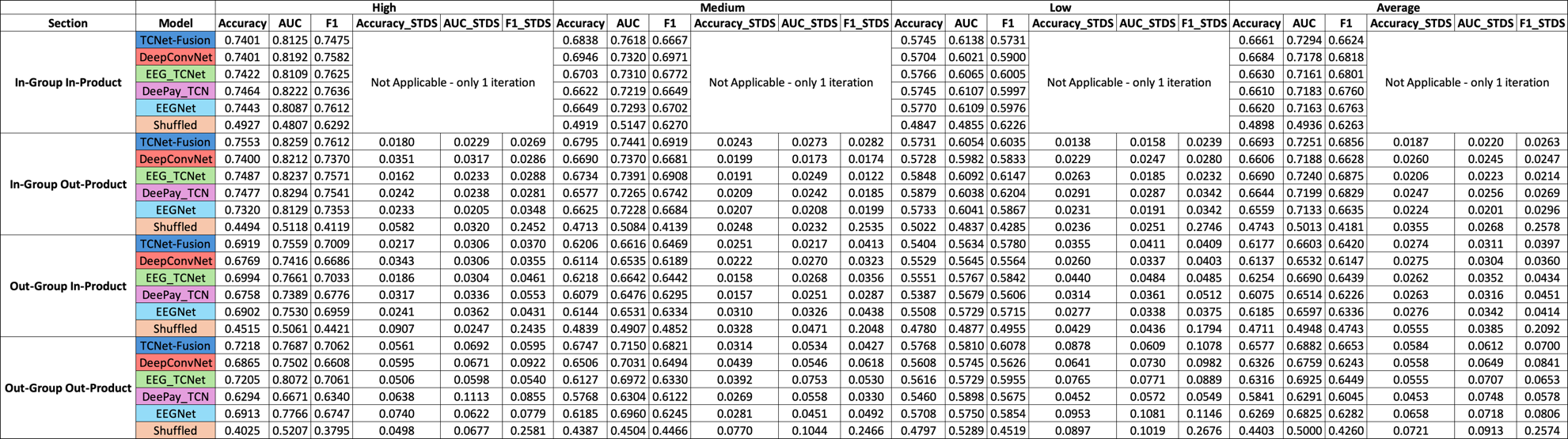

**Table W16**. Neural network’s prediction results for Part 1 WTP. The different models are coded by color – TCNet-Fusion in light blue, DeepConvNet in red, EEGTCNet in green, DeePay in dark blue, EEGNet in purple and shuffled DeePay in orange. The table is organized in 4 main rows – each for each prediction attempt – In-Group In-Product – we used all participants and all products randomly for the training, validation and testing separations. In-Group Out-Product – the test set consisted of products the network did not train on. Out-Group In-Product – the test set consisted of participants the network did not train on. Out-Group Out-Product – the test set consisted of both participants and products the network did not train on. Between the columns you can find the different performance measures – Accuracy, AUC and F1 for classification. You can also find their STDS between the columns. The different data parsing is also in the column names when 20_80 is the High separation with the top 20% and bottom 20% of the data; 20_35_65_80 is the Medium separation with 20-35% and 65-80% of the data, and 50 is the low separation with values between 35-50% and 50-65%.

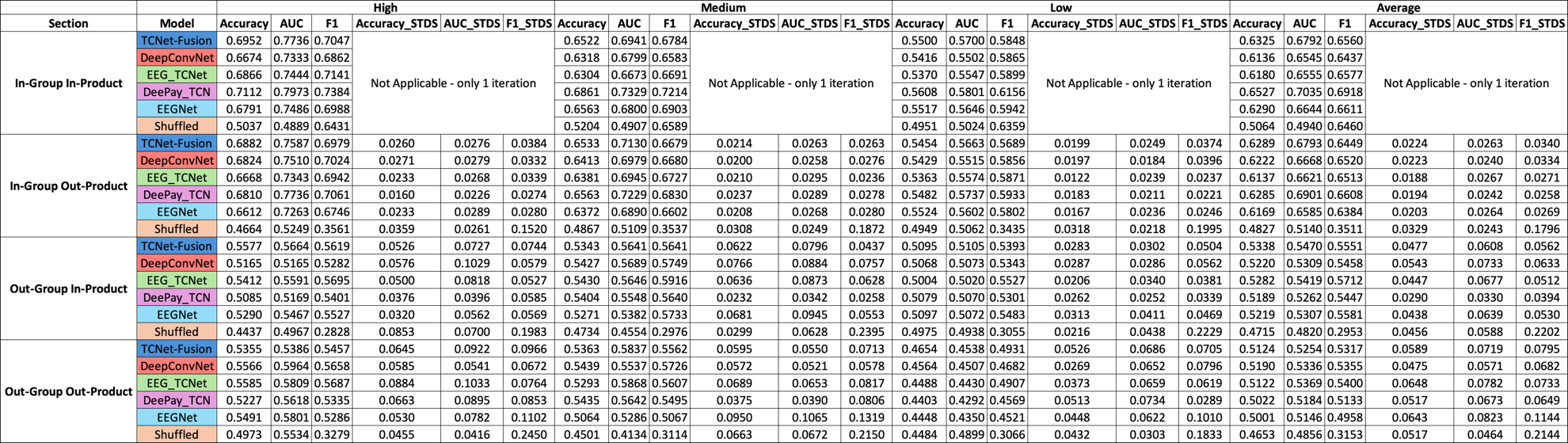

**Table W17.** Neural network’s results for Part 2 WTP prediction. The different models are coded by color – TCNet-Fusion in light blue, DeepConvNet in red, EEGTCNet in green, DeePay in dark blue, EEGNet in purple and shuffled DeePay in orange. The table is organized in 4 main rows – each for each prediction attempt – In-Group In-Product – we used all participants and all products randomly for the training, validation and testing separations. In-Group Out-Product – the test set consisted of products the network did not train on. Out-Group In-Product – the test set consisted of participants the network did not train on. Out-Group Out-Product – the test set consisted of both participants and products the network did not train on. Between the columns you can find the different performance measures – Accuracy, AUC and F1 for classification. You can also find their STDS between the columns. The different data parsing is also in the column names when 20_80 is the High separation with the top 20% and bottom 20% of the data; 20_35_65_80 is the Medium separation with 20-35% and 65-80% of the data, and 50 is the low separation with values between 35-50% and 50-65%.

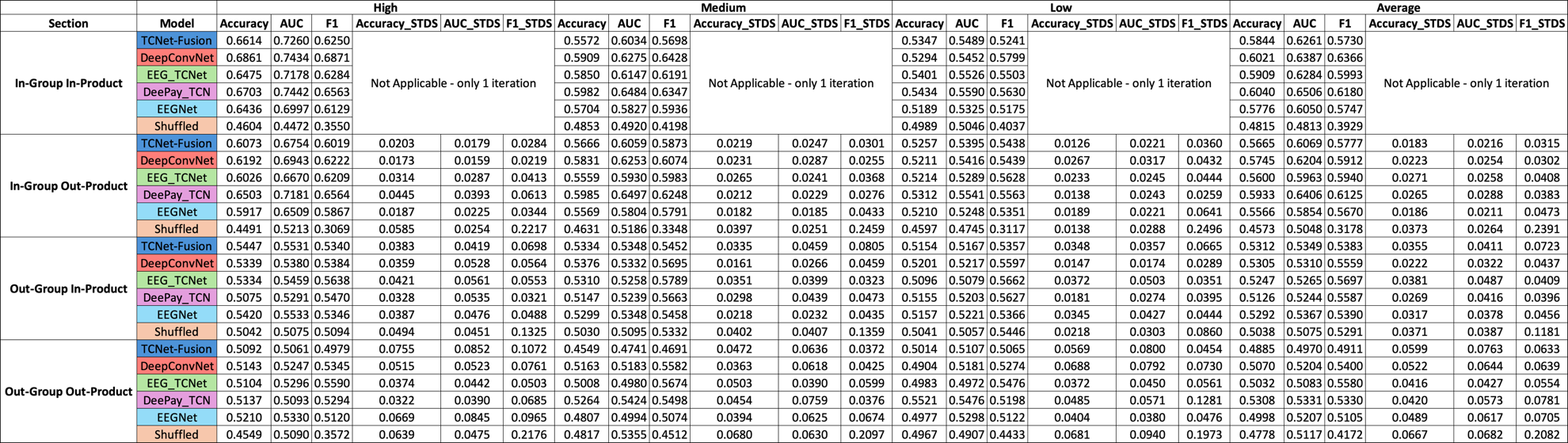

**Table W18.** Neural network’s results for Part 2 Liking prediction. The different models are coded by color – TCNet-Fusion in light blue, DeepConvNet in red, EEGTCNet in green, DeePay in dark blue, EEGNet in purple and shuffled DeePay in orange. The table is organized in 4 main rows – each for each prediction attempt – In-Group In-Product – we used all participants and all products randomly for the training, validation and testing separations. In-Group Out-Product – the test set consisted of products the network did not train on. Out-Group In-Product – the test set consisted of participants the network did not train on. Out-Group Out-Product – the test set consisted of both participants and products the network did not train on. Between the columns you can find the different performance measures – Accuracy, AUC and F1 for classification. You can also find their STDS between the columns. The different data parsing is also in the column names when 20_80 is the High separation with the top 20% and bottom 20% of the data; 20_35_65_80 is the Medium separation with 20-35% and 65-80% of the data, and 50 is the low separation with values between 35-50% and 50-65%.

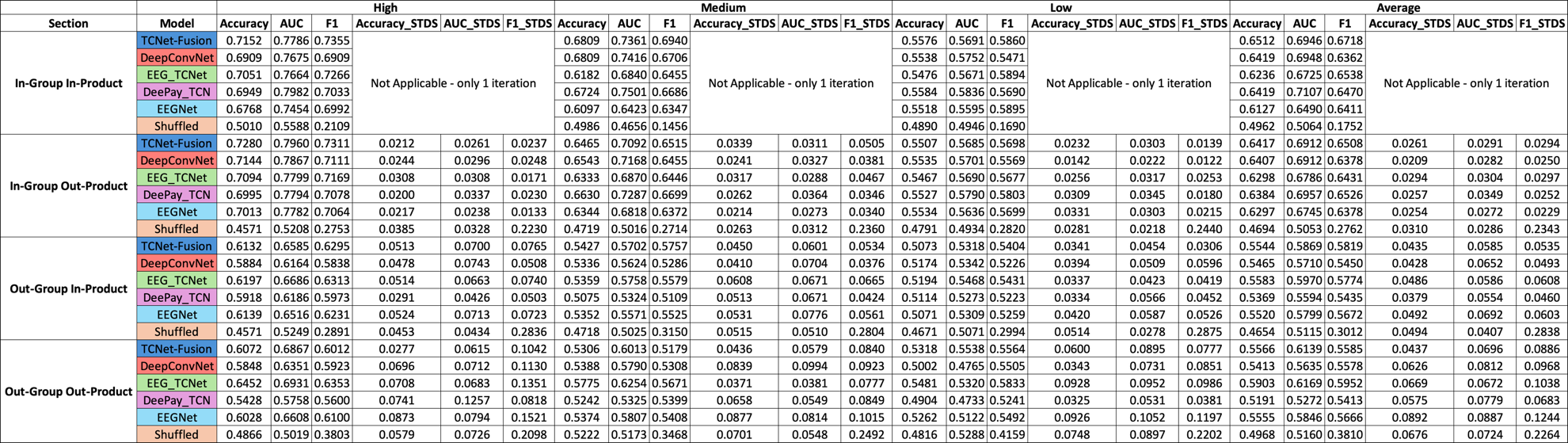
**Table W19**. Neural network’s results for Part 3 WTP prediction with Part 1 EEG signal. The different models are coded by color – TCNet-Fusion in light blue, DeepConvNet in red, EEGTCNet in green, DeePay in dark blue, EEGNet in purple and shuffled DeePay in orange. The table is organized in 4 main rows – each for each prediction attempt – In-Group In-Product – we used all participants and all products randomly for the training, validation and testing separations. In-Group Out-Product – the test set consisted of products the network did not train on. Out-Group In-Product – the test set consisted of participants the network did not train on. Out-Group Out-Product – the test set consisted of both participants and products the network did not train on. Between the columns you can find the different performance measures – Accuracy, AUC and F1 for classification. You can also find their STDS between the columns. The different data parsing is also in the column names when 20_80 is the High separation with the top 20% and bottom 20% of the data; 20_35_65_80 is the Medium separation with 20-35% and 65-80% of the data, and 50 is the low separation with values between 35-50% and 50-65%.

### **Web Appendix G: Logistic Regression**

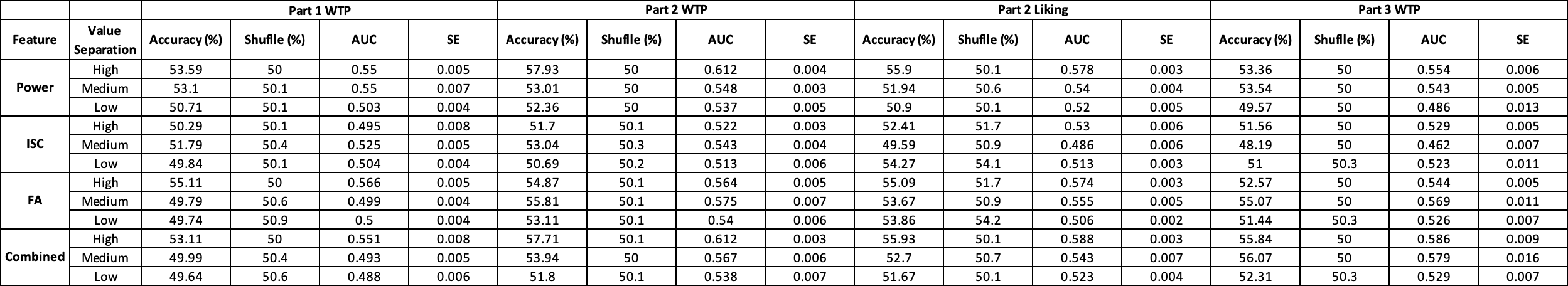

**Table W20**. Logistic-regression benchmark results across EEG feature types, experimental parts, and value-separation levels. This table summarizes the performance of the logistic-regression classifiers used as linear benchmarks for the EEG-based prediction tasks. Rows correspond to the four feature sets evaluated: frequency-band power, inter-subject correlation (ISC), frontal asymmetry (FA), and a combined model with all features (Combined). For each feature set, results are shown separately for the *High*, *Medium*, and *Low* value-separation levels used in the main analysis. The columns are organized into four groups representing the different prediction targets: Part 1 WTP, Part 2 WTP, Part 2 Liking, and Part 3 WTP (delayed valuation using Part 1 EEG to predict Part 3 behavior). Within each group, we report averaged Test Accuracy (%), Shuffled label chance level (%), AUC, and accuracy SE. Overall, the table shows that logistic-regression models achieved near-chance accuracy across most feature sets, separation levels, and parts, with only small and inconsistent above-chance effects at *High* (and occasionally *Medium*) separation-particularly for frequency-band power and FA. ISC and Combined-feature models rarely exceeded chance, and all feature types converged to chance at *Low* separation. These results provide a linear baseline for comparison with the neural-network models reported in the main text.

### **Web Appendix H: Neural Networks Trial Distribution**

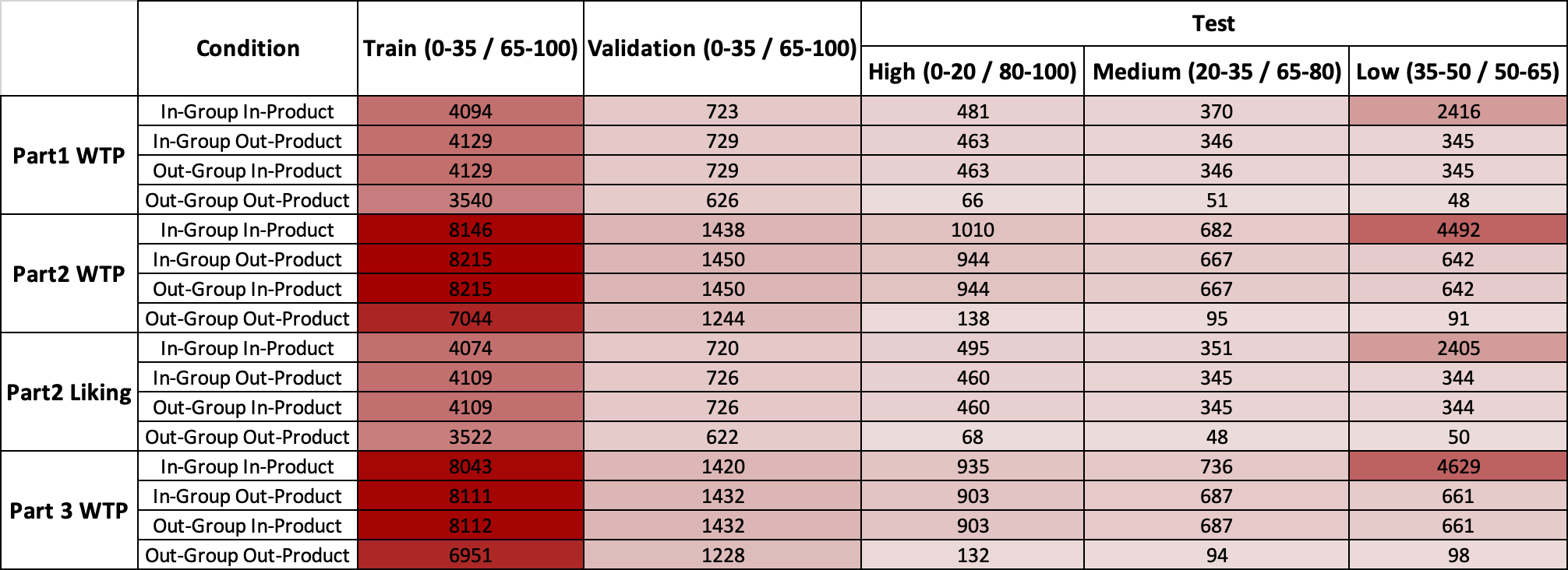

**Table W21**. **Distribution of trial counts across experimental conditions and dataset splits.** The table presents the number of trials for each condition, separated by experimental part (Part 1 WTP, Part 2 WTP, Part 2 Liking, Part 3 WTP – delayed valuation using Part 1 EEG to predict Part 3 behavior), group membership (in-group vs. out-group), and product type (in-product vs. out-product), across data splits used for modeling. Columns correspond to the training set (0 – 35% / 65% – 100%), validation set (0 – 35% / 65% – 100%), and three test subsets defined by value ranges: high (0 – 20% / 80% – 100%), medium (20% – 35% / 65% – 85%), and low (35% – 50% / 50% – 65%). Cell colors reflect trial counts using a red gradient, with darker shades indicating higher counts.
